# A multi-strategy antimicrobial discovery approach reveals new ways to combat *Chlamydia*

**DOI:** 10.1101/2023.11.30.569351

**Authors:** Magnus Ölander, Daniel Rea Vázquez, Karsten Meier, Lieke Mooij, Johanna Fredlund, Fabiola Puértolas-Balint, Eduard Calpe, María Rayón Díaz, Karin van der Wal, Bjoern O. Schroeder, Barbara Susanne Sixt

**Affiliations:** Department of Molecular Biology, Umeå University, Umeå, Sweden; The Laboratory for Molecular Infection Medicine Sweden (MIMS), Umeå University, Umeå, Sweden; Umeå Centre for Microbial Research (UCMR), Umeå University, Umeå, Sweden

**Keywords:** intracellular bacteria, drug discovery, phenotypic screening, virtual screening, small molecules, narrow-spectrum antibiotics, antibiotic synergies, antibiotic mode of action, bacterial persistence, autophagy

## Abstract

While the excessive use of broad-spectrum antibiotics is a major driver of the global antibiotic resistance crisis, more selective therapies remain unavailable for the majority of bacterial pathogens. This includes *Chlamydia* spp., which cause millions of urogenital, ocular, and respiratory infections each year. We here report major conceptual additions to the available toolkit for antichlamydial discovery, including both experimental and computational approaches. Moreover, we report the to date most comprehensive search of the chemical space for novel antichlamydial activities, which identified over sixty compounds that are chemically diverse, structurally different from known antibiotics, non-toxic to human cells, and highly potent in blocking *Chlamydia* growth. While some compounds caused a reversible block in *Chlamydia* development, others could eradicate both established and persistent infections in a bactericidal manner. The most potent antichlamydials displayed compelling selectivity, some also synergies with clinically used antibiotics, as well as interactions profiles enabling predictions of molecular modes of action. Moreover, one compound displayed reduced antichlamydial efficacy in autophagy-deficient cells, suggesting a host-targeted activity. Altogether, we suggest that these novel antichlamydials could serve as tools for advancing our understanding of *Chlamydia* biology and as chemical starting points for developing more sustainable therapeutics for one of the most successful groups of intracellular pathogens.

## Introduction

Antimicrobial resistance is a major societal challenge that is quickly aggravating and estimated to cause over 10 million deaths annually by 2050 ^1^. A prime driver of this alarming development is the heavy use of broad-spectrum antibiotics that select for resistance not only in the targeted pathogen but in a wide range of exposed microbes ^2^. By perturbing commensal and environmental microbial communities, these broad-acting drugs can damage human health and ecosystems as well ^3,4^. Clearly, there is an urgent need to develop more selective and sustainable treatment alternatives, especially for those pathogens that account for significant global consumption of antibiotics.

A prime example is *Chlamydia trachomatis*, the world’s most common bacterial agent of sexually transmitted infections (STIs), responsible for over 130 million cases annually ^5^. *Chlamydia* STIs can cause serious complications and long-term sequelae, such as pelvic inflammatory disease, infertility, and pregnancy complications ^6^. *C. trachomatis* also causes trachoma, a devastating blinding ocular disease ^7^. Moreover, closely related *Chlamydia* spp. have significant medical and veterinary impact as well ^8,9^.

Lacking effective vaccines, *C. trachomatis* is fought by antibiotic treatment ^10^, which in the context of STIs must include treatment of sexual contacts ^6^, while mass treatment of entire communities remains common practice in managing trachoma ^11^. This sums up to substantial global antibiotic use for the control of this single pathogen alone. Even worse, since selective therapeutics are unavailable, current treatment relies entirely on broad-acting antibiotics, in particular doxycycline and azithromycin ^10^. Yet, these antibiotics are especially disruptive to the human microbiota ^12^, and their use in anti-*Chlamydia* therapy indeed exacerbates resistance development in bystander pathogens ^13–15^. At the same time, treatment failure rates of up to 5-23% have been reported for *Chlamydia* STIs ^16^.

*Chlamydia* spp. are obligate intracellular bacteria that must invade the cells of their host to cause disease. The bacteria then thrive inside membrane-enclosed vacuoles, called inclusions, engage in highly specific host-pathogen interactions, and undergo complex developmental transitions ^17^. More specifically, the infectious form of the bacteria, the elementary body (EB), invades a host cell and then differentiates into the reticulate body (RB), the replicative form, which multiplies within the inclusion. Eventually, RBs differentiate back into EBs, which are released by host cell lysis or extrusion of the inclusion about 48-72 hours post infection (hpi) ^18^. This strong dependence on host cells should make *Chlamydia* spp. well suited for developing narrow-spectrum therapeutics that can target *Chlamydia* or its interactions with host cells specifically without causing collateral damage. Yet, the development of such has not yet been successful.

To develop selective antichlamydial therapies, we need suitable molecular targets and chemical scaffolds acting on those. Because *Chlamydia*-host interactions remain ill-understood at the molecular level, we chose to approach this need by using a target-agnostic approach, which combined both experimental and virtual screening of large libraries of small drug-like molecules, followed by an in-depth analysis of candidate molecules by determinations of compound potency, toxicity, selectivity, interactions, and mode of interference with *Chlamydia* development. As a result, we here report the identification of numerous potent selective antichlamydials that may serve both as chemical starting points for developing novel therapeutics and as tools to identify therapeutically targetable host-pathogen interactions specific to *Chlamydia*.

## Results

### Development of a screening assay for novel chemical inhibitors of *C. trachomatis* growth

To efficiently screen for antichlamydial compounds, we developed a screening assay that operates in a 384- well plate format, requires minimal manual handling of plates, and can monitor, in parallel, inhibition of intracellular bacterial growth and compound toxicity towards host cells. Specifically, we used a GFP-expressing strain of *C. trachomatis* L2/434/Bu (CTL2-GFP) to infer bacterial growth from bulk GFP fluorescence, and we used the metabolic conversion of resazurin into its product resorufin to infer host cell viability from bulk resorufin fluorescence (Fig S1A). In brief, human cervical epithelial (HeLa) cells were infected with CTL2-GFP in suspension and seeded into plates containing compounds in dimethyl sulfoxide (DMSO). After 26 hours of incubation, resazurin was added, followed by further incubation for 2.5 hours, and measurement of GFP and resorufin fluorescence. Wells treated solely with DMSO served as negative control, while wells treated with ciprofloxacin or staurosporine served as positive controls for bacterial growth inhibition and cytotoxicity, respectively.

During assay optimization, we found a seeding density of 4,000 cells per well and an infection dose of 30 inclusion-forming units (IFUs) per cell to be ideal, as this gave near-maximal resorufin fluorescence without signal saturation (Fig S1B) and high GFP fluorescence without cytotoxicity (Fig S1C-E). Importantly, we found that CTL2-GFP and HeLa cells showed reasonable tolerance to DMSO, up to concentrations of around 0.4% and 1.6%, respectively (Fig S1F). Moreover, in an interleaved-signal plate layout ^19^, both readouts resulted in data that were robust and uniform throughout the plate, with Z’ factors of 0.53-0.62 and 0.51-0.73 for bacterial growth inhibition and toxicity, respectively (Fig S1G-H).

We benchmarked our assay using a set of traditional antibiotics, all of which gave dose-dependent reductions in GFP fluorescence with minimal effects on host cell viability (Fig S2A). Minimum inhibitory concentrations (MICs) calculated from these data spanned more than five orders of magnitude and were, with one exception, within or close to the ranges reported ^20–30^ (Fig S2B).

Taken together, we developed an assay with excellent performance parameters that can reliably measure the inhibition of *C. trachomatis* growth over a wide range of compound concentrations.

### Identification of novel antichlamydials through experimental compound library screening

We screened a library of 36,785 small molecules displaying drug-like properties and covering a wide chemical space (Fig 1A). In this process, we also complemented the assay described above with an imaging-based readout (Fig 1B). After measurements of bulk GFP and resorufin fluorescence, cells were fixed, stained with the DNA dye Hoechst, and imaged at a high-content imaging platform to detect chlamydial inclusions (GFP) and host cell nuclei (Hoechst). The total area and number of inclusions in individual wells and the number of nuclei then served as additional proxies for chlamydial growth and host cell viability.

**Fig 1.**
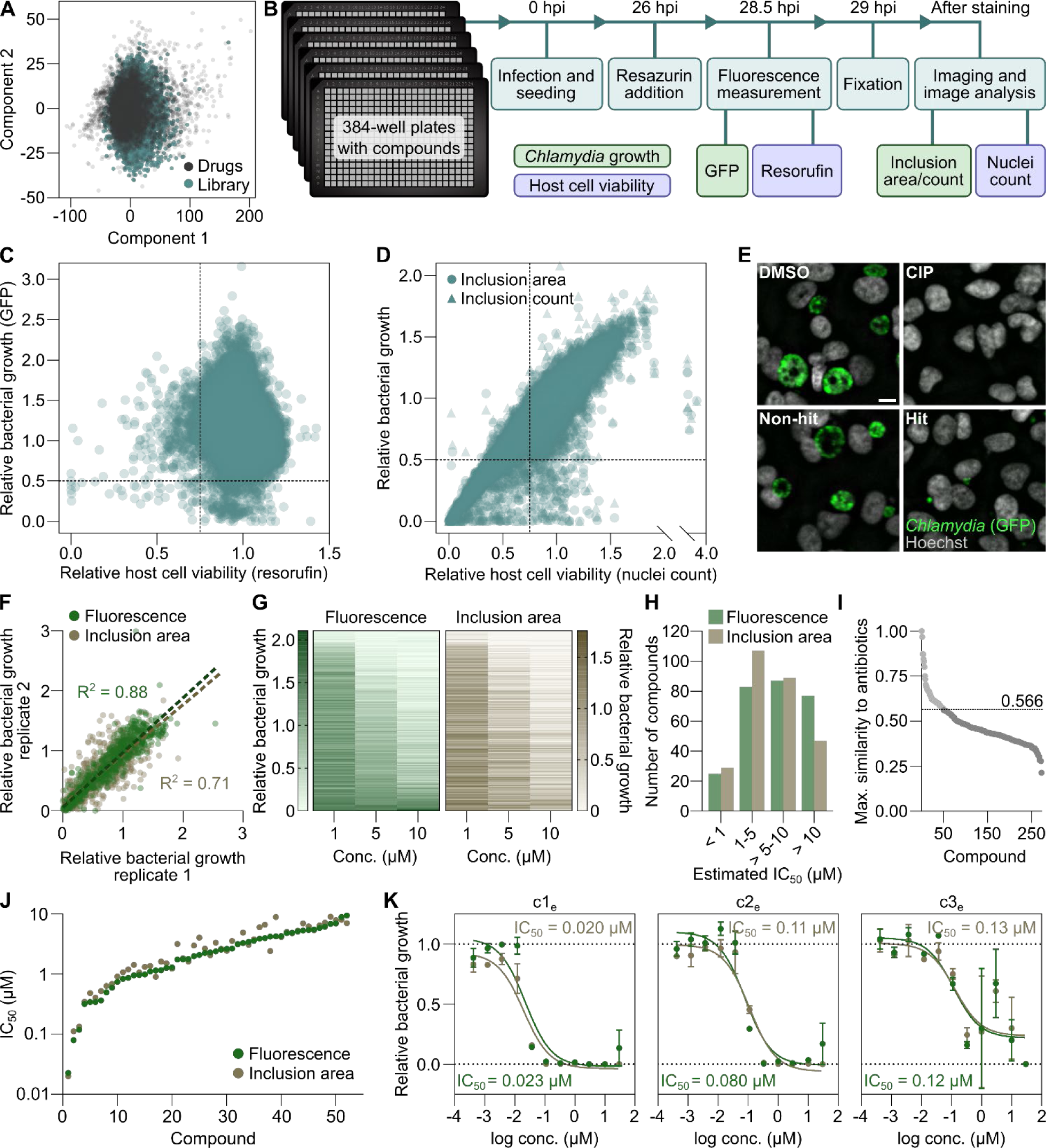
**Identification of novel antichlamydials through experimental compound library screening**. **(A)** Chemical diversity of the screening library of 36,785 compounds, illustrated by principal component analysis based on molecular descriptors calculated with PaDEL-Descriptor ^31^. ‘Drugs’ denotes 6,798 compounds from the Drug Repurposing Hub ^32^, included to represent the chemical space of pharmaceutical drugs. **(B)** Screening protocol, highlighting major experimental steps, time points, and measurements. Compounds were tested at 10 µM. **(C)** Bulk fluorescence measurements of GFP and resorufin, representing *C. trachomatis* growth and host cell viability, respectively. The dashed lines indicate the cut-offs used for hit selection. **(D)** Image-based measurements of inclusion area and count and host cell nuclei count, representing *C. trachomatis* growth and host cell viability, respectively. The dashed lines indicate the cut- offs used for hit selection. **(E)** Representative image examples of a non-hit and a hit, as well as positive (ciprofloxacin, CIP) and negative (DMSO vehicle) controls for bacterial growth inhibition. Scale bar is 10 µm. **(F)** The 271 hits from the screen retested in duplicate with the screening assay protocol at 1, 5, and 10 µM, highlighting the overall consistency between the two replicates. **(G)** Bacterial growth inhibition achieved by the 271 hit compounds, shown at the single-compound level. The compounds are sorted by mean bulk fluorescence across the three concentrations. **(H)** Estimated IC_50_ of the 271 hits. **(I)** Maximal structural similarity, represented by Tanimoto coefficients, between the hits and 506 known antibiotics in the Drug Repurposing Hub ^32^. The dashed line indicates the maximum value of compounds prioritized. **(J)** Potency of the 52 priority compounds, determined with the screening assay protocol (mean of n = 3). **(K)** Dose-response curves of the three most potent compounds identified by experimental screening (c1_e_-c3_e_) (mean ± SD, n = 3). Lines indicate curve fits used for IC_50_ calculation.

When analyzing the data derived from bulk fluorescence measurements in the screen, we identified 303 compounds that displayed antichlamydial activity (> 50% reduction in GFP fluorescence) but were non- toxic towards the host cells (< 25% reduction in resorufin fluorescence) (Fig 1C and Table S1). Using the same cut-offs in the analysis of image-derived data, we identified 136 compounds as non-toxic antichlamydials according to the number of nuclei and either total inclusion area or inclusion count (Fig 1D- E and Table S1). We then generated a decision tree to integrate data from both fluorescence measurements and imaging for a final classification into hits and non-hits and selected a total of 271 compounds for further evaluation (Fig S3 and Table S1).

A retesting of the 271 compounds in duplicate at three concentrations confirmed 179 compounds to be antichlamydial and non-toxic (> 50% reduction in GFP fluorescence and inclusion area, and < 25% reduction in resorufin fluorescence, at ≤ 10 µM; Fig 1F-G and Table S2). We further estimated compound potencies, using as measure the half-maximal inhibitory concentration (IC_50_), considered less ambiguous than MICs ^33^ (Fig 1H and Table S2), and decided to prioritize compounds with estimated IC_50_ ≤ 5 µM (according to both GFP fluorescence and inclusion area) for further follow-up. Moreover, because our objective was to discover antichlamydials with novel chemical structures, not yet associated with antimicrobial activities, we excluded compounds that resembled known antibiotics (Fig 1I and Table S2), resulting in a list of 52 priority compounds (Fig S4 and Table S2).

Subsequently, we retested these 52 compounds in triplicate at a wide range of concentrations to enable more accurate potency determinations (Fig S5, and Table S3). Ten compounds showed submicromolar IC_50_ values for both growth parameters (*i.e.*, GFP fluorescence and inclusion area) (Fig 1J and Table S3). Three were particularly potent, with IC_50_ values around or below 0.1 µM (Fig 1K).

Altogether, our efforts led to the identification of a set of highly potent non-toxic antichlamydial compounds that are structurally different from known antibiotics.

### Predictive modeling enabling antichlamydial discovery through virtual screening

As a complement to our experimental approach, we took advantage of our data to develop a machine-learning-based prediction model, which then enabled us to identify additional compounds with potential antichlamydial activity through virtual screening (Fig 2A).

**Fig 2.**
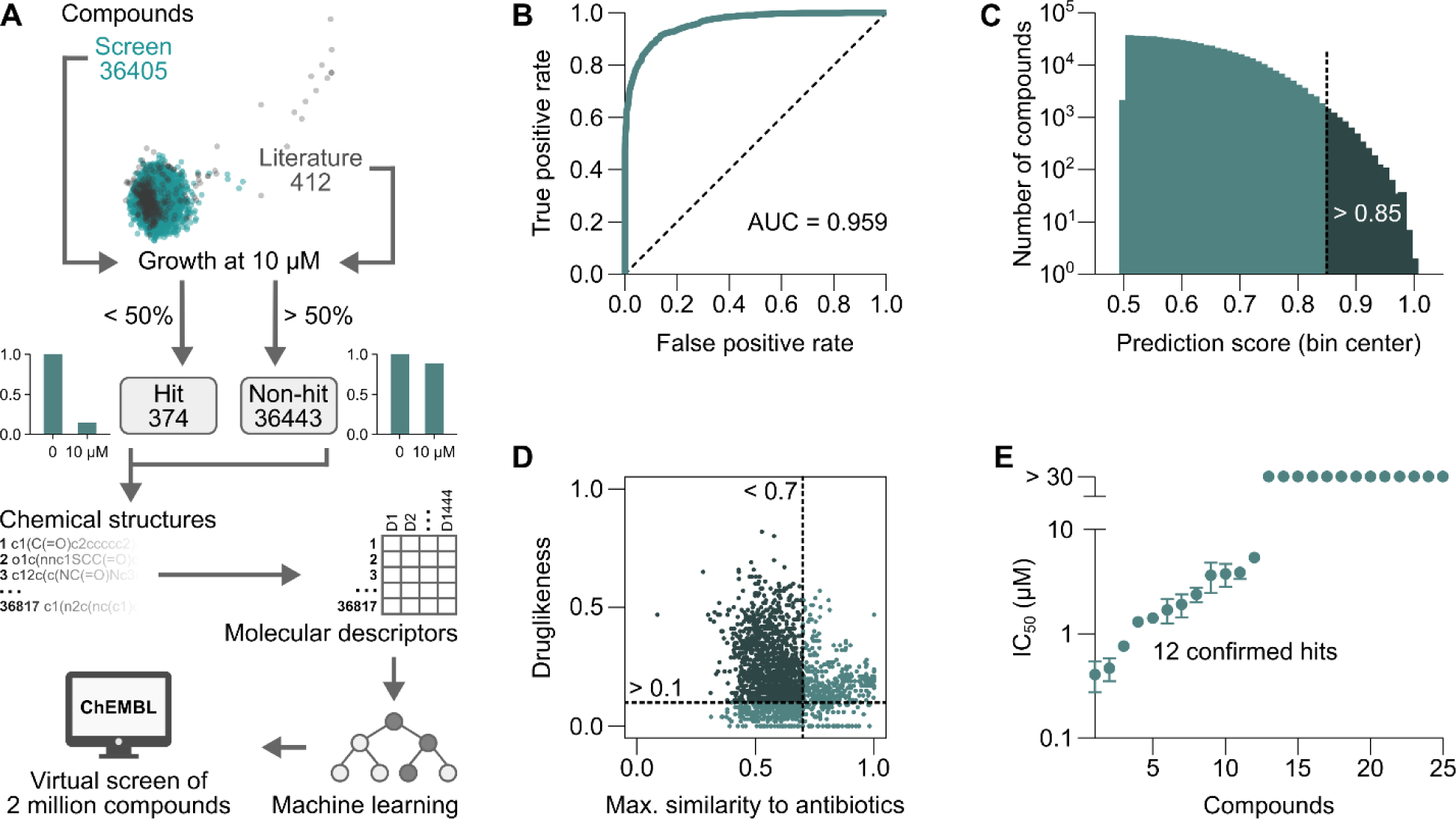
Predictive modeling enabling antichlamydial discovery through virtual screening. (A) Schematic illustration of model development. **(B)** ROC-AUC plot showing the performance of the model after training. **(C)** Histogram of predicted antichlamydial hits from a virtual screen of the ChEMBL database. The dashed line indicates the cut-off used to filter the hits based on prediction scores. **(D)** Maximal structural similarity to known antibiotics (Tanimoto coefficients) and quantitative estimate of druglikeness (QED) of the filtered hits. The dashed lines indicate the cut-offs used for further filtering. **(E)** Potency of 25 compounds subjected to experimental testing, as measured by bulk GFP fluorescence using the screening assay protocol (mean ± SD, n = 3).

The model was based on our own dataset (including all 36,405 compounds from the experimentally screened library for which structures were available) and 412 additional compounds previously reported to have been tested for antichlamydial activity (Table S4). All molecules in this expanded dataset were classified as hits or non-hits using the same criterion as used in our experimental screen (> 50% growth inhibition at 10 µM). Using the PaDEL-Descriptor tool ^31^, we then calculated 1,444 1D and 2D molecular descriptors for all compounds. Subsequently, these were used as input to train a random forest classifier, which, given a new structure, predicts a molecule’s probability of anti-chlamydial activity. The model achieved an area under the receiver operating characteristic curve (ROC-AUC) of 0.959 on the training data (Fig 2B), with 88.9% correctly classified instances (based on ten-fold cross-validation).

Next, we applied our model to perform predictions on the ChEMBL database, which includes over two million chemically diverse drug-like compounds ^34^. This virtual screen resulted in 5,474 possible antichlamydials with prediction scores > 0.85 (Fig 2C). Filtering these to retain only one structural isomer per compound reduced the number to 2,871. We further retained only such that had a low structural similarity to known antibiotics (Tanimoto coefficient < 0.7) and a quantitative estimate of druglikeness (QED ^35^) > 0.1 (Fig 2D), which left 174 compounds (Table S5). After determining their commercial availability, we selected 25 affordable compounds for experimental validation. Twelve of these (representing two series of chemical analogs and one more distinct molecule) indeed showed antichlamydial activity with IC_50_ values below 10 µM (Fig 2E and Table S6), with the top three displaying submicromolar IC_50_ values. These three compounds share a sulfonamide core structure with previously identified antichlamydials included in the training set ^36^, yet have relatively low overall structural similarity (Tanimoto coefficients of around 0.63-0.78) to these previously described antichlamydials.

Taken together, we present here a predictive model that could successfully identify highly potent antichlamydials through virtual screening of a very large compound database.

### Machine learning-aided discovery of persistence-inducing compounds

Certain antibiotics, in particular beta-lactams, induce “chlamydial persistence”, a stress response often characterized by the formation of aberrant bodies (ABs), *i.e.*, highly enlarged RBs ^37^. These persistent RBs do not divide nor differentiate into EBs but can survive within their host cell for prolonged periods of time. Upon removal of the stressor, chlamydial replication and development can resume ^37^. While persistence inducers may not be the preferred starting points for the development of novel clinical drugs, they may serve as tools for deciphering the mechanistic basis of this ill-understood phenomenon.

To search our experimental screening data for potential persistence-inducing compounds, we again enlisted the assistance of machine learning to help identify images displaying AB morphologies in inclusions.

Two different binary classification models were trained on a wide range of features of chlamydial inclusions calculated by image analysis. During model training, we used images of infected cells treated with beta-lactam antibiotics as ‘hits’, and images of DMSO controls as ‘non-hits’. We then applied the trained models to all images from the screen, resulting in 504 compounds predicted as hits by at least one model. Manual inspection showed that only two of the compounds (persistence-inducing compounds 1-2, in brief c1_p_-c2_p_) indeed induced AB formation (Fig 3A-B). Yet, when inspecting images of 504 randomly selected non-hits, none showed signs of AB formation.

**Fig 3.**
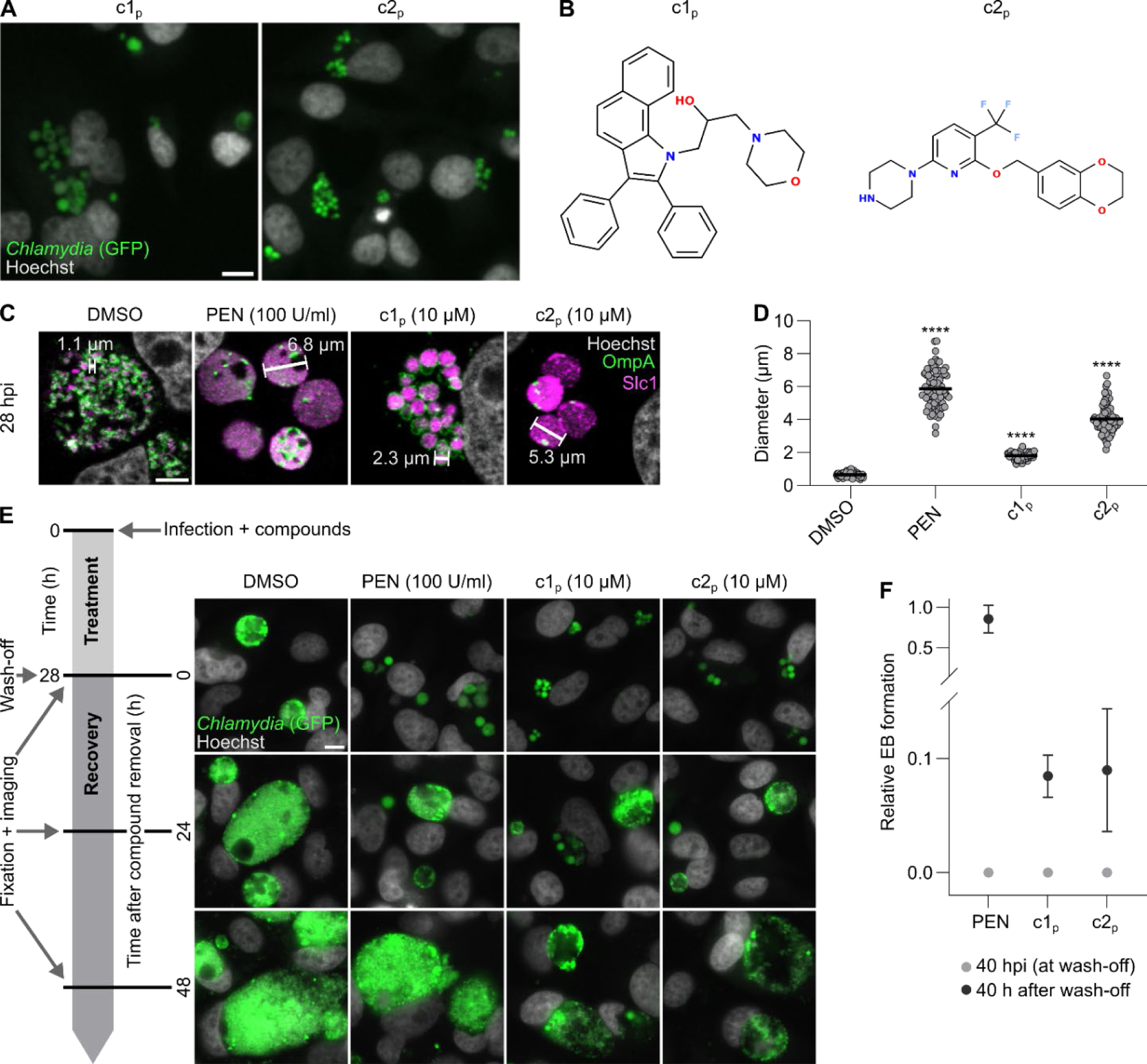
Machine learning-aided discovery of persistence-inducing compounds. (A) Persistence-inducing compounds identified by predictive modelling applied to images from the experimental compound library screen. Scale bar is 10 µm. **(B)** Structures of the identified compounds. **(C-D)** Confocal microscopic confirmation of a persistent phenotype (formation of ABs) in HeLa cells infected with CTL2 and exposed to the indicated compounds. (C) Representative images of one of three experiments. Bacteria were detected using antibodies specific for OmpA (the major outer membrane protein) and Slc1 (a type III secretion chaperone). The annotated lines show the diameter of representative individual bacteria. Scale bar is 5 µm. (D) Size of compound-treated bacteria (collated data of n = 3, line at mean, 30 bacteria measured per condition and experiment, two-way ANOVA with Dunnett’s multiple comparisons test). **(E)** Recovery of the growth of CTL2-GFP in HeLa cells after compound removal. The images are representative of three experiments. Scale bar is 10 µm. **(F)** EB formation by CTL2-GFP and its recovery from persistence in HeLa cells after exposure to the compounds (10 µM) or penicillin G (100 U/ml) for 40 h (mean ± SD, n = 3). Data was normalized to values from DMSO-treated wells at wash-off. PEN, penicillin G.

We then used confocal microscopy to study compound effects on bacterial morphology at higher resolution (Fig 3C-D). As expected, treatment with penicillin G caused enlarged bacteria (*i.e.*, ABs) with diameters of around 4-8 µm to form. AB formation was also observed in cells treated with c1_p_ or c2_p_, with mean bacterial diameters of about 2 µm and 4 µm, respectively, well within the reported range for ABs ^38^. We further observed that persistence induction by c1_p_ and c2_p_ was reversible, as ABs disappeared and bacterial growth resumed, when the compounds were washed out after 28 h of treatment (Fig 3E). Moreover, c1_p_ and c2_p_ blocked the formation of EBs and this blockage was also reversible (Fig 3F).

Collectively, these data demonstrate that the image-based aspects of our screening approach can also be used to identify compounds causing more complex morphological phenotypes, typified here by the discovery of two novel persistence inducers.

### Determination of antibacterial selectivity of our best antichlamydials

To verify that the antichlamydial activities described above were not restricted to HeLa cells, we selected top compounds from the experimental (c1_e_-c5_e_) and virtual (c1_v_-c5_v_) screens and tested them against CTL2- GFP in cells of different origins. Beside HeLa cells, we chose A2EN cells, as they are human endocervical epithelial cells derived from a non-cancerous tissue _39_. Moreover, we included cells from animal species, *i.e.*, monkey (Vero), mouse (BALB/3T3), guinea pig (JH4), and chicken (UMNSAH), that may in part serve as hosts in future *in vivo* testing. Overall, we observed IC_50_ values to be similar across all cell lines for c1_e_, c2_e_, and c4_e_, while c3_e_ and c5_e_ showed some variability (Fig 4A and S6A). Compounds c1_v_-c5_v_ displayed similar potency in HeLa and BALB/3T3 cells (Fig 4A and S6B). Critically, c1_e_, c2_e_, c4_e_, and c1_v_-c5_v_ were non-toxic up to the highest tested concentration (3 or 30 µM; Fig 4B and S7A-B). While, at 30 µM, c3_e_ and c5_e_ were toxic to some cell lines, both were well tolerated at lower potently antichlamydial concentrations.

**Fig 4.**
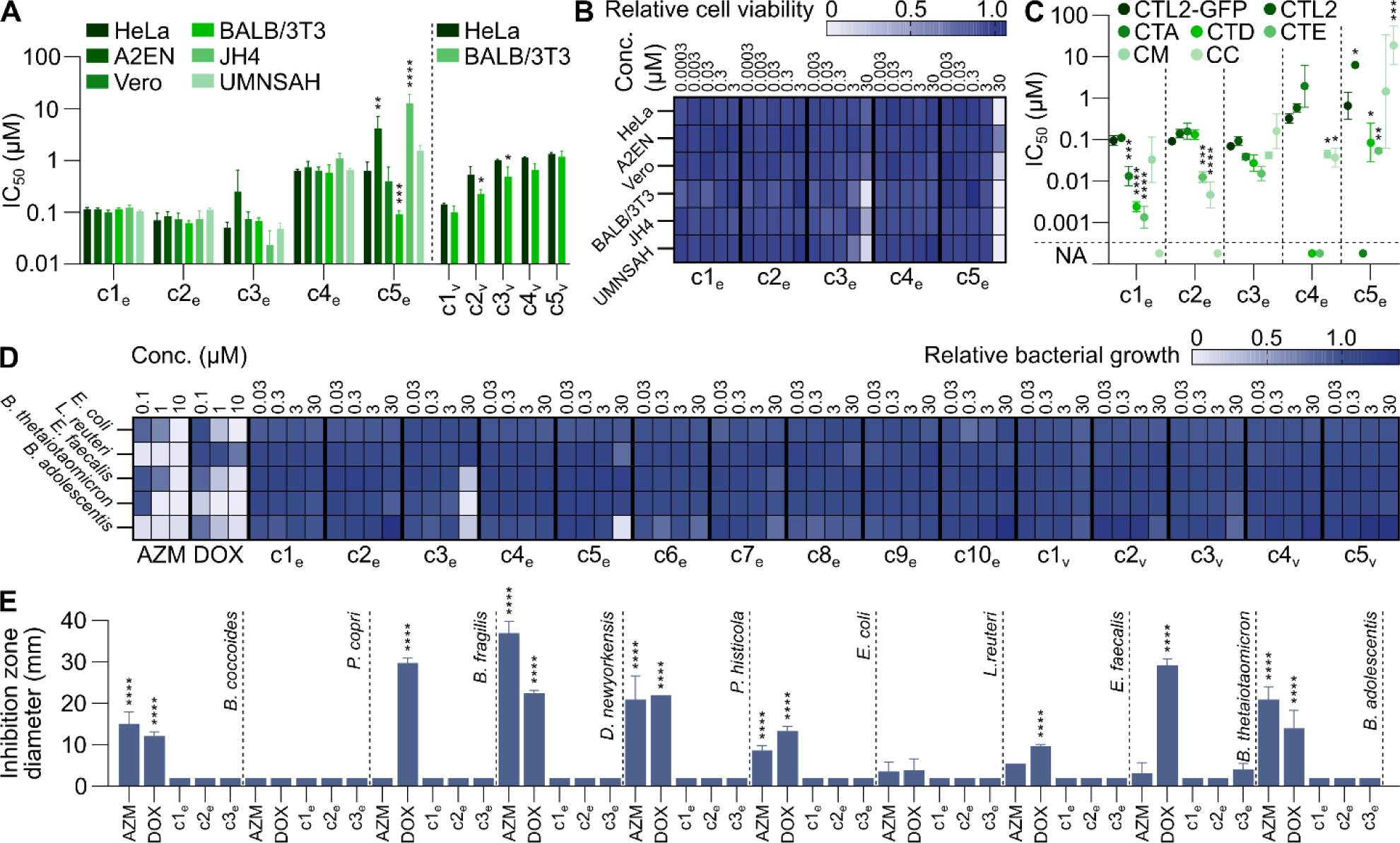
Determination of antibacterial selectivity of our best antichlamydials. (A) Potency of selected top compounds in different cell lines infected with CTL2-GFP, as measured by bulk GFP fluorescence (mean ± SD, n = 3, two-way ANOVA with Sidak’s multiple comparisons test of HeLa vs. other cell lines). **(B)** Effects of selected top compounds on viability in different cell lines, measured in parallel with (A), using resorufin fluorescence (mean of n = 3). **(C)** Potency of selected top compounds in HeLa cells infected with the indicated strains, as measured by inclusion area (mean ± SD, n = 3, two-way ANOVA with Dunnett’s multiple comparisons test vs. CTL2-GFP). NA, not active at the concentrations tested. **(D)** Effect of top compounds on the growth of five gut microbiota species in liquid medium, monitored by OD_600_ measurements (mean of n = 3). **(E)** Effect of c1_e_-c3_e_ on the growth of additional gut bacteria species in radial diffusion assays (mean ± SD, n = 2-5, one-way ANOVA with Dunnett’s multiple comparisons test vs. non-inhibited DMSO-treated controls). AZM, azithromycin; DOX, doxycycline.

Next, we evaluated the effect of c1_e_-c5_e_ on other strains and species of *Chlamydia*. IC_50_ values for c1_e_- c4_e_ differed by less than twofold between CTL2-GFP and parental CTL2, and those seen for c5_e_ by less than tenfold (Fig 4C). With the exception of c5_e_, all compounds were active against the trachoma serovar, *C. trachomatis* A/HAR-13 (CTA). Moreover, c1_e_-c3_e_ and c5_e_ showed similar or higher potency against the genital serovars, *C. trachomatis* D/UW-3/Cx (CTD) and *C. trachomatis* E/Bour (CTE), than against CTL2. Surprisingly, c4_e_ was inactive against these strains at all concentrations tested. All compounds also showed similar or higher potency against *Chlamydia muridarum* (CM) than against CTL2, while results for *Chlamydia caviae* (CC) were variable with c3_e_-c5_e_ being active and c1_e_-c2_e_ being inactive at all tested concentrations. As our objective was to discover compounds that inhibit *Chlamydia* selectively but spare commensal bacteria, we further tested fifteen of our top compounds (c1_e_-c10_e_ and c1_v_-c5_v_) against five bacterial species representing the four dominating phyla of the human gut microbiota _40_. In general, the compounds did not affect the growth of the bacteria in liquid media at up to 30 µM (Fig 4D and S8). Exceptions were c3_e_ and c5_e_, which at 30 µM were inhibitory to some species. In stark contrast, azithromycin fully inhibited the growth of all tested species at 10 µM or lower, and doxycycline inhibited all but *L. reuteri* (Fig 4D), already previously reported to resist the action of this drug ^41^. Of note, c1_e_-c3_e_ had little or no effect on the growth of an even larger set of species tested in radial diffusion assays, in which the bacteria were exposed to compounds for 24-48 h before measurements of growth inhibition zones (Fig 4E).

Taken together, our top compounds are potent and selective inhibitors of the intracellular growth of *Chlamydia* spp. and do not, in general, affect commensal bacteria of the human gut microbiota.

### Determination of between-compound interactions and interactions with clinical antibiotics

Next, we set out to explore the potential for synergistic interactions among our top compounds and between them and antibiotics currently used in the clinics to treat *Chlamydia* or other infectious diseases.

To establish a framework for testing combination treatments, we tested twelve clinical antibiotics in all possible pairwise combinations against CTL2-GFP in HeLa cells. Among the 66 combinations, we observed instances of all interaction types (synergistic, antagonistic, and additive) (Fig 5A). We then tested our best compounds (c1_e_-c10_e_ and c1_v_-c5_v_) in combination with the twelve antibiotics and found that c4_e_ and c9_e_ showed strong or moderate synergistic interactions with some of the antibiotics (Fig 5B and S9). Most prominently, c4_e_ synergized strongly with doxycycline and ofloxacin. Finally, we tested c1_e_-c3_e_ in pairwise combinations with c1_e_-c10_e_ and c1_v_-c5_v_ and observed relatively strong synergistic interactions between c1_e_ and c4_e_ and between c2_e_ and c6_e_ (Fig 5B and S9).

**Fig 5.**
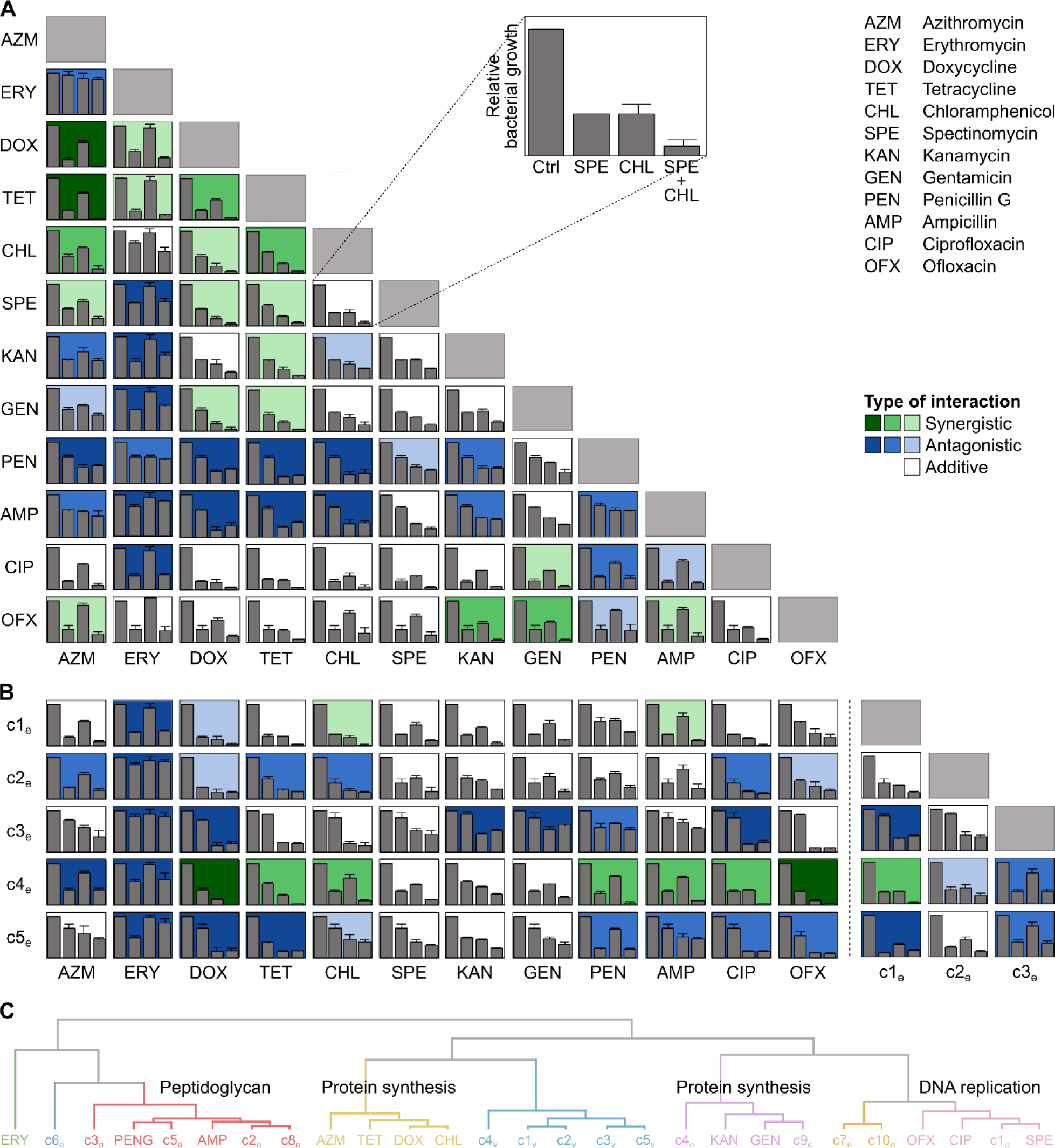
Determination of between-compound interactions and interactions with clinical antibiotics. (A) Bacterial growth inhibition in HeLa cells infected with CTL2-GFP and classification of interaction type for all pairwise combinations of twelve clinical antibiotics, tested at IC_50_ (mean ± SD, n = 3). Interaction type was classified based on calculations of epistasis, as previously described ^42^. Darker colors indicate stronger synergistic or antagonistic interactions. **(B)** Bacterial growth inhibition and classification of interaction type for pairwise combinations of selected compounds and antibiotics, tested at IC_50_ as in (A) (mean ± SD, n = 3). **(C)** Hierarchical clustering based on principal component analysis of pairwise interaction profiles (first three principal components), using the calculated epistasis values. The naming of the clusters is based on the targets of the known antibiotics they contain.

The kind of overall pairwise interaction profiles generated here have previously been used to cluster antibiotics by mode of action (MoA) and to identify compounds with novel mechanisms, for instance in *E. coli* _42_. Thus, we used hierarchical clustering based on principal component analysis to compare our interaction profiles (Fig 5C). Antibiotics with similar MoA indeed formed clusters, supporting the validity of the approach. For instance, the ribosome-targeting antibiotics azithromycin, chloramphenicol, doxycycline, and tetracycline formed one cluster, while the peptidoglycan-targeting antibiotics ampicillin and penicillin G formed another, also including c2_e_, c3_e_, c5_e_, and c8_e_ (Fig 5C). Moreover, c1_e_ clustered with the fluoroquinolones ciprofloxacin and ofloxacin, which interfere with bacterial DNA replication, while compounds c4_e_ and c9_e_ clustered with the aminoglycosides gentamicin and kanamycin, which disturb proofreading during translation. Unsurprisingly, considering their structural similarities, c1_v_-c5_v_ clustered tightly together, indicating that they act in a similar way.

Overall, these interaction data provide first hints towards potential MoA of c4_e_ and c9_e_ and suggest that these compounds (or such with similar MoA) may have a potential to be used in combination therapies.

### Determining the stage in *Chlamydia* development targeted by our best antichlamydials

A key event in the chlamydial developmental cycle is the generation of EBs that can exit the host cell to spread the infection (Fig 6A). Because our compounds can prevent intracellular replication of *C. trachomatis*, and EB formation is a late event that occurs once RBs have ceased to replicate, we were not surprised to see that c1_e_-c5_e_ and c1_v_-c5_v_ abrogated EB formation by CTL2-GFP in HeLa cells (Fig 6B). To further investigate which stage of intracellular growth was interfered with, we added compounds (c1_e_-c10_e_) at a later time point and then assessed bacterial growth at 48 hpi, based on bulk GFP fluorescence.

**Fig 6.**
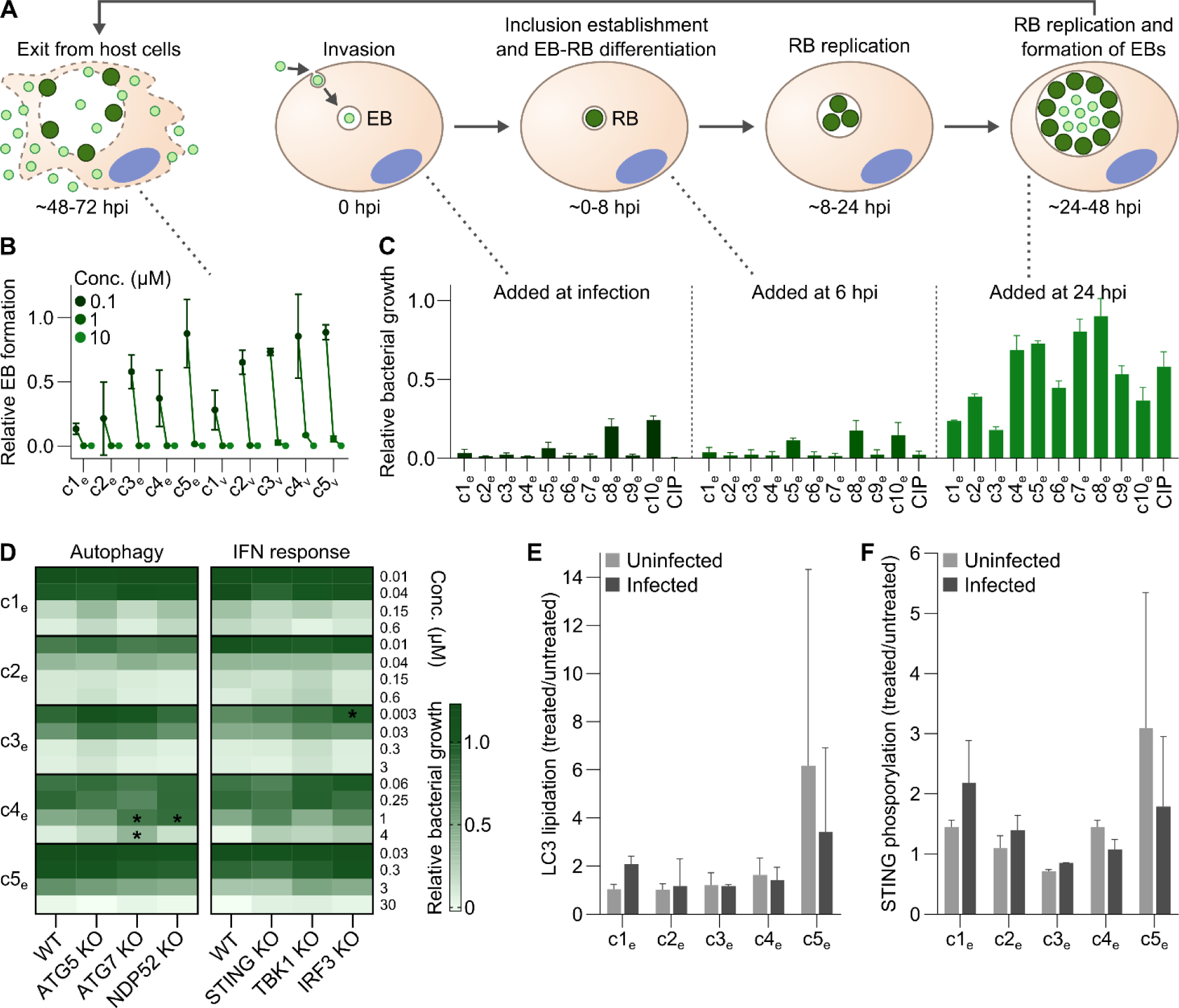
Determining the stage in *Chlamydia* development targeted by our best antichlamydials. (A) Illustration of the chlamydial developmental cycle **(B)** Effect of top compounds on EB formation in HeLa cells infected with CTL2-GFP (mean ± SD, n = 3). **(C)** Growth inhibition after addition of top compounds (10 µM) at different points in the developmental cycle, tested in HeLa cells infected with CTL2-GFP, as measured at 48 hpi by bulk GFP fluorescence (mean ± SD, n = 3). **(D)** Antichlamydial activity of our best compounds in A2EN cells (wild-type (WT) or mutant as indicated) infected with CTL2-GFP, as measured at 28.5 hpi by bulk GFP fluorescence (mean of n = 3, two-way ANOVA with Dunnett’s multiple comparisons test). **(E-F)** LC3 lipidation (E) and STING phosphorylation (F) after compound exposure in uninfected A2EN cells and cells infected with CTL2, represented as the ratio of compound-treated and DMSO-control samples (mean ± SD, n = 3, two-way ANOVA with Sidak’s multiple comparisons test). CIP, ciprofloxacin.

Compound addition at 6 hpi (after invasion and early inclusion formation) resulted in essentially identical growth inhibition compared to addition at the time of infection (Fig 6C). As expected, addition at 24 hpi (to an established infection) reduced bacterial growth less strongly, though most compounds still caused significant levels of inhibition. Notably, c1_e_-c3_e_ reduced growth still by around 60-80% (Fig 6C), clearly demonstrating that they act on the replicative phase of the infection.

In the replicative phase, the ability of *C. trachomatis* to thrive intracellularly strongly depends on its ability to evade the intrinsic defenses of its host cell ^43^. Hence, we reasoned that some of our antichlamydials may act by boosting the defenses or by weakening bacterial evasion. As a first foray into this question, we tested the activity of c1_e_-c5_e_ in A2EN cells deficient for key proteins of either the autophagic machinery (ATG5, ATG7, NDP52) or the STING pathway of the type I IFN response (STING, TBK1, IRF3) (Fig S10). In most cases, we observed no significant differences between wild-type and defense-defective cells (Fig 6D). However, c4_e_ was less effective in cells deficient for ATG7 or NDP52 (a trend also observed for ATG5), suggesting that a functional autophagy machinery may be necessary for its full antichlamydial activity. We further analyzed the induction of the defense programs in treated cultures by measuring LC3 lipidation and STING phosphorylation (Fig 6E-F). We observed that c5_e_ displayed a tendency to induce both, but highly variable and in infected as well as in uninfected cells, likely connected to its higher toxic potential (Fig 4B). We further noticed a tendency, albeit also not statistically significant, for an infection-specific induction of both programs by c1_e_. Interestingly, c4_e_ did not induce overall enhanced levels of LC3 lipidation.

In conclusion, the top antichlamydials identified in this study affect the bacteria in their replicative RB phase. As a consequence, they also abrogate formation of EBs and can thus stall the spread of infection. Moreover, we found that autophagy may be involved in the action of some of the compounds.

### Compounds can eradicate both established and persistent infections in a bactericidal manner

Since our best compounds displayed inhibitory effects even when added to established infections (Fig 6C), we wanted to assess whether a longer treatment duration would lead to total eradication. Thus, we infected HeLa cells with CTL2-GFP, added compounds (c1_e_-c5_e_) at 24 hpi, and replenished them at 48 and 72 hpi, followed by wash-off at 96 hpi and incubation until 168 hpi. Bacterial growth was assessed every 24 h, based on bulk GFP fluorescence. At sufficient concentrations, all compounds acted bactericidal, as they blocked further growth during the course of treatment and did not allow growth to resume after wash-off (Fig 7A). We saw similar trends with azithromycin and doxycycline (Fig 7B), which are known to exert bactericidal effects on *C. trachomatis* ^44^. Moreover, imaging supported these findings (Fig 7C).

**Fig 7.**
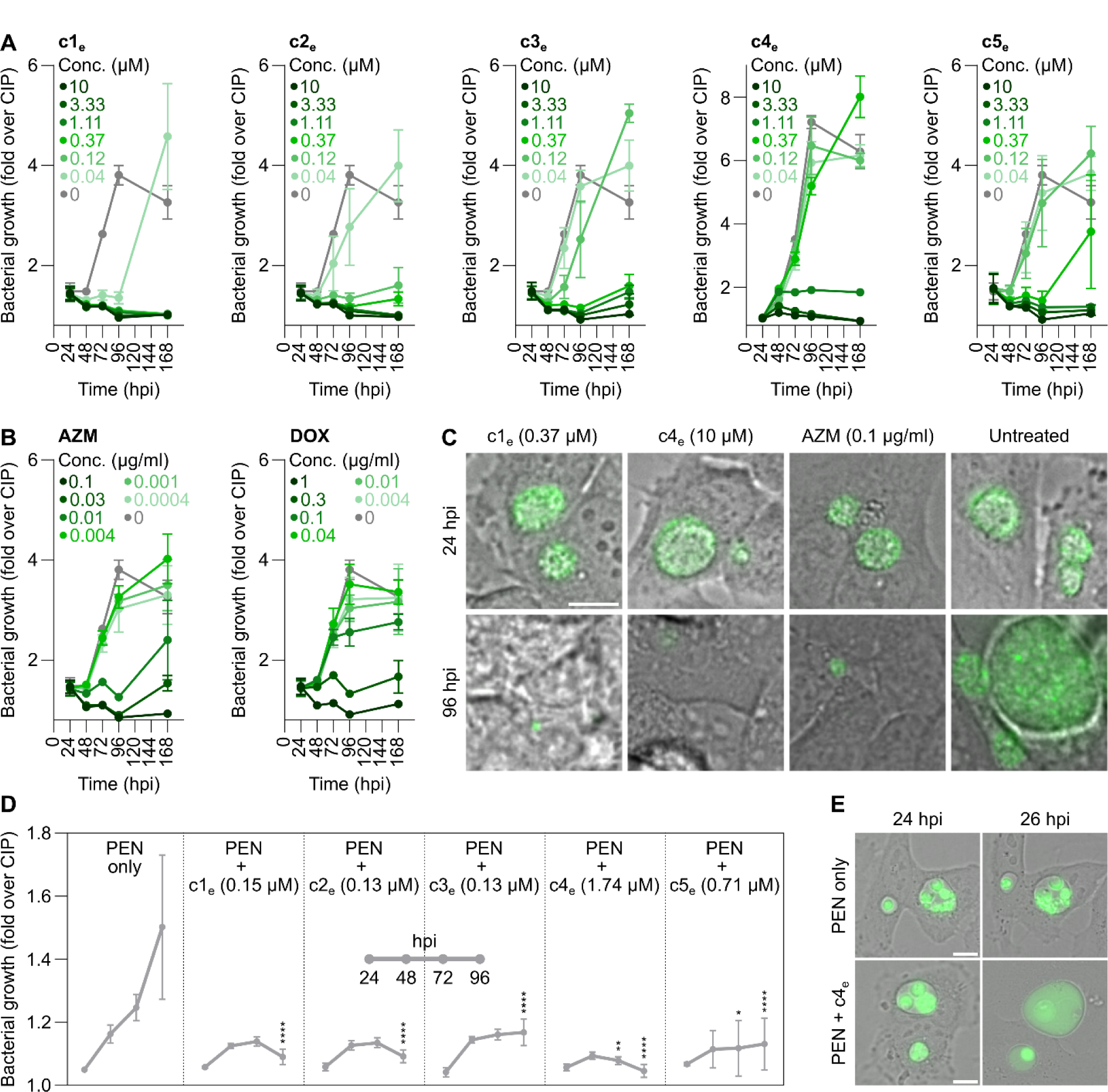
Compounds can eradicate both established and persistent infections in a bactericidal manner. (A-B) Time course of bacterial growth in HeLa cells infected with CTL2-GFP and treated with different concentrations of c1_e_-c5_e_ (A) or azithromycin or doxycycline (B) (mean ± SD, n = 3). Compounds and antibiotics were first added at 24 hpi, then replenished daily until wash-off at 96 hpi. The data was normalized to ciprofloxacin-treated wells, thus showing fold-increase of GFP fluorescence above background levels. (C) Images corresponding to selected time points and concentrations in (A) and (B). Scale bar is 10 µm. (D) Time course of bacterial growth in HeLa cells infected with CTL2-GFP in the presence of penicillin G (present throughout the experiment) and treated with c1_e_-c5_e_ from 24 hpi with daily replenishment (mean ± SD, n = 3, two-way ANOVA with Dunnett’s multiple comparisons test of compound vs. penicillin G at each time point). **(E)** Live imaging of cells treated as in (D). Scale bar is 10 µm. AZM, azithromycin; DOX, doxycycline; PEN, penicillin G.

Next, we wanted to evaluate if our compounds would be effective against persistent infections. Thus, we performed a similar time-course experiment but adding penicillin G at infection to induce persistence. In cells treated with penicillin G alone, we observed a slow but steady increase in fluorescence throughout the experiment, due to continuous enlargement of ABs. Yet, in the presence of compounds (c1_e_-c5_e_, added at 24 hpi and replenished daily), there was almost no increase, and several compounds reduced fluorescence to close to background by 96 hpi (Fig 7D). Moreover, in live imaging of compound-treated persistent infections, we observed a noticeable effect of c4_e_ as early as two hours post-treatment. The number of ABs decreased, and inclusions began to fill with diffuse GFP fluorescence, indicating lysis of ABs (Fig 7E).

In summary, these results indicate that our top antichlamydials have bactericidal activity against *C. trachomatis* and can eradicate established as well as persistent infections.

## Discussion

Here, we reported major conceptual additions to the available toolkit for antichlamydial discovery, a significant expansion of the pool of known antichlamydial molecules, and the identification and in-depth characterization of a set of highly potent and selective antichlamydials, which can serve as tools and may inspire the development of novel therapeutics. In brief, we (a) developed a simple, robust, and information-rich assay for high-throughput screening for non-toxic inhibitors of *Chlamydia* growth (Fig S1 and S2); (b) applied this assay for a comprehensive search of the chemical space of drug-like small molecules (Fig 1 and S3); (c) integrated new and previously reported data on antichlamydials into a machine-learning based model that can predict antichlamydial activities *in silico* (Fig 2); and (d) applied this model in a virtual search to probe an even larger chemical space (Fig 2). Combined, these efforts led to the identification of over sixty potent antichlamydials that are chemically diverse and dissimilar to known antibiotics (Fig 1-2). In-depth characterization of top molecules revealed that they (with exceptions) were: (a) active in all tested cell lines (Fig 4A); (b) non-toxic to host cells at concentrations 30-fold higher than their IC_50_ (Fig 4B); (c) active against a range of *Chlamydia* strains and species (Fig 4C); (d) selective, *i.e.*, not growth-inhibitory towards microbiota species (Fig 4D-E); (e) able to prevent inclusion establishment and EB formation when present at the beginning of the infection (Fig 6B-C); (f) able to eradicate established inclusions in a bactericidal manner (Fig 7A); and (g) able to eradicate persistent infections (Fig 7D). Some compounds also showed synergistic interactions with each other and/or clinically used antibiotics (Fig 5B and S9).

To date, antichlamydial discovery strategies have been primarily based on the compound-first principle, involving phenotypic screens for molecules blocking pathogen growth *^e.g.^*^, 36,45–47^, to be later followed by MoA determinations. A few studies described target-based approaches, which involved experimental screening of compound libraries for molecules inhibiting specific targets ^48,49^ or virtual screening for molecules that bind to selected target structures ^50,51^. However, target-based approaches require a profound knowledge of the target, such as its structure, molecular function, and biological role ^52^, hence their applicability to *Chlamydia* remains restricted by our limited understanding of its biology. Moreover, while target-based approaches may uncover molecules that can inactivate their target potently *in vitro*, these may lack other properties needed for activity in a biological system, such as the ability to penetrate the multiple membrane barriers that shield intracellular bacteria ^53^.

Phenotypic high-throughput screening of drug libraries requires a simple, fast, and cost-effective set- up. Yet, *Chlamydia*’s lifestyle entails a need for detecting intracellular pathogen growth, which makes screening protocols more demanding and explains why the majority of previous screens, with exceptions *^e.g.,^* ^36,46^, involved rather small compound selections. When compared to previously applied strategies in antichlamydial discovery *^e.g.^*^, 36,45,46,54–60^, our assay stands out by combining maximum simplicity with cost-effectiveness, and an ability to provide a wealth of information for each compound (Fig 1B). By seeding already pre-infected cells in compound-containing plates, we bypassed many steps carried out individually in most described protocols. Moreover, by omitting medium exchange or wash steps prior to the fluorescence measurements, we significantly lowered the risk for (cross-)contamination or wash-off effects. Moreover, the additional recording of imaging data made the screen more robust, as visual information can help to identify false positives and to exclude compounds with non-desired MoA.

Because experimental screening will always remain time- and resource-consuming, no matter how simple the assay, an intriguing idea is to exploit available information to first define what properties makes a molecule likely to be active and then to use this information for a virtual pre-screening. Along this line, Karhu et al. mapped nineteen known antichlamydial compounds into the chemical space, based on their physicochemical properties, and then compared these with a library of 502 molecules to select only such with properties most close to the known antichlamydials for experimental testing ^61^. More powerful virtual screening for novel antimicrobials can be enabled by training machine-learning-based prediction models on data from compound library screens together with descriptors of chemical structures ^62^. We took a similar approach, using data taken from our own experimental screen and the literature to train a binary random forest classifier that can predict the antichlamydial activity of input structures (Fig 2). Along that way, we encountered hurdles, such as the problem of class imbalance ^63^, which required us to employ class balancing techniques to improve model performance, and difficulties in comparing data between studies that employed a plethora of different readouts. These problems may be mitigated in the future by generating an expanded in-house dataset for model training. Nevertheless, the present model enabled screening many more compounds than otherwise feasible, and thereby the, to our knowledge, first comprehensive target-agnostic virtual screen ever performed for *Chlamydia* spp.

The reasons for a significant number of *Chlamydia* infections to become chronic or recurrent or to prove otherwise recalcitrant to treatment are not fully understood. However, *Chlamydia* persistence could be a major contributing factor ^37^. Persistence can be induced by various stress conditions. This includes nutrient deprivation ^64,65^ and exposure to cytokines ^66^, but also exposure to antibiotics, most notably beta-lactams ^67,68^. Doxycycline and azithromycin may also induce persistence when used at for the bacteria sub-lethal doses ^69,70^. Significantly, once in a persistent state, the bacteria are less susceptible to eradication by antibiotic treatment ^44,71–73^. Hence, effective therapeutics would be expected to not be prone to induce persistence and to be able to eradicate persistent inclusions. In this context, it was reassuring to see that our top compounds did not induce morphological signs of persistence and were able to eradicate, in a bactericidal manner, established (Fig 7A) and persistent inclusions (Fig 7D). However, we also identified two new persistence-inducing compounds (Fig 3), which may be valuable tools for deciphering the molecular basis of this ill-understood phenomenon.

For novel antichlamydial therapeutics to be more sustainable than the currently available, they would also need to be more selective (narrow spectrum), though a broad activity against *Chlamydia* spp. may still be desirable. Indeed, our top compounds were active against a range of *Chlamydia* serovars and species (Fig 4C), suggesting that they could inspire new treatments for various *Chlamydia* diseases, potentially also zoonotic and veterinary infections. Critically, when tested against a representative selection of gut microbes, our top compounds displayed striking selectivity, in stark contrast to the broad action of doxycycline and azithromycin (Fig 4D-E). Yet, ultimately, both efficacy and selectivity will also need to be confirmed *in vivo*. Understanding compound MoA could deepen our knowledge of *Chlamydia* biology, identify targets for future target-based discovery approaches, and guide the further chemical optimization of the molecules. In this context, our observations in regards to compound interactions might provide interesting hypotheses to be tested, as antibiotics with similar MoA often share similar interactions profiles ^42^. For instance, we observed that in relation to its interaction profile, c1_e_ clustered with fluoroquinolones (Fig 5C), suggesting that c1_e_ may interfere with DNA replication in *Chlamydia*. Moreover, c4_e_ and c9_e_ clustered with aminoglycosides (Fig 5C), suggesting that c4_e_ and c9_e_ may also compromise ribosome (proofreading) function in *Chlamydia*. The latter would also be consistent with the bacteriolytic activity of c4_e_ (Fig 7E), as aminoglycosides are known to lyse bacterial cells ^74,75^. Given the highly selective action of the compounds, it also seems plausible that some might interfere with more specific features of *Chlamydia* development or host-pathogen interactions. Our observation that c4_e_ was less effective in autophagy-deficient cells (Fig 6D), while not causing a bulk stress-induced increase in autophagy (Fig 6E), may for instance suggest that this compound sensitizes the pathogen to host cell-autonomous defenses.

Taken together, our innovative multi-strategy antimicrobial discovery approach enabled this to date most comprehensive search of the chemical space for novel *Chlamydia* growth-inhibitory small molecules, which identified more than sixty promising antichlamydials. In the future, these molecules can serve as tools to transform our understanding of *Chlamydia* pathogenesis, but also as chemical starting points for the development of more sustainable therapeutics, which are urgently needed for this medically challenging and prevalent group of bacterial pathogens.

## Significance

The impact of this work has two major dimensions, as we here provide strategies and tools for addressing both a major crisis in medicine and pressing questions in the field of host-pathogen interactions. The alarming rise in antimicrobial resistance is a serious threat to modern medicine. This problem needs to be tackled at its root, by reducing the widespread unnecessary exposure to broad-acting drugs. In this context, *Chlamydia* spp. represent a significant case, accounting for massive global use of broad-spectrum antibiotics, with both treatment failures and resistance-promoting effects on bystander pathogens being well-documented. In the present work, we developed and implemented new antichlamydial discovery tools, including a simple, effective, and informative assay for experimental screening, applicable to this challenging intracellular pathogen, as well as the first computational model for *in silico* prediction of antichlamydial activities from chemical structures. Combining both, we conducted the to date most comprehensive search of the chemical space of drug-like small molecules for antichlamydial activities, and thereby identified over sixty novel antimicrobials that are chemically diverse and structurally different from known antibiotics. We further conducted in-depth analyses of compound potency, toxicity, selectivity, interactions, and mode of interference with *Chlamydia* development. Together, these efforts significantly advance the field towards providing more selective therapeutics for *Chlamydia* spp. At the same time, *Chlamydia* spp. have a unique developmental biology and are important models for dissecting mechanisms of intracellular virulence and host-pathogen interactions, and thereby also for shedding light on the biology of human cells. Yet, our molecular understanding is sparse because molecular genetic manipulation of the pathogen remains challenging. Hence, by identifying molecules that can potently but selectively perturb the intracellular growth, host-pathogen interactions, or development of *Chlamydia*, this work promotes the identification of its unique biological features and virulence strategies.

## Materials and Methods

### Cell culture

HeLa (ATCC CCL-2), Vero (ATCC CCL-81), HEK293T (ATCC CRL-3216), BALB/3T3 (ATCC CCL-163), and UMNSAH/DF-1 (ATCC CRL-12203) cells were cultivated in Dulbecco’s Modified Eagle’s Medium (DMEM; Gibco) supplemented with 10% heat-inactivated fetal bovine serum (FBS; Gibco). In experiments involving bulk fluorescence measurements or live cell imaging, a phenol red-free DMEM was used to reduce background. A2EN cells ^39^ were cultivated in Keratinocyte-SFM (Gibco) and JH4 cells (ATCC CCL-158) in Ham’s F-12 (Kaighn’s) medium (Gibco), both supplemented with 10% heat-inactivated FBS. In experiments including fluorescence measurements in A2EN and JH4 cells, their specific media were replaced by phenol red-free DMEM 2.5 hours before the measurements (*i.e.*, prior to resazurin addition). All cell cultures were maintained at 37°C and 5% CO2 in a humidified incubator.

### Generation of STING pathway- and autophagy-deficient cell lines

Cell lines deficient for the indicated genes were generated using CRISPR/Cas9-mediated genome editing ^76^. A lentiviral delivery system was used to introduce genes coding for Cas9 nuclease and gene specific sgRNAs. The following sgRNAs, previously validated by Sanjana et al ^77^, were cloned into vector lentiCRISPRv2 ^77^ (Addgene 52961):ATG5 (5’- AGATCAAATAGCAAACCAAT-3’, 5’-TTCCATGAGTTTCCGATTGA-3’), ATG7 (5’- AGAAGAAGCTGAACGAGTAT-3’, 5’-TAGGGTCCATACATTCACTG-3’), NDP52 (5’-CAACAAATCAGCTAAACAGC-3’, 5’-AATCAGAGTGGATCAGCTTC-3’), STING (5’-GGATGTTCAGTGCCTGCGAG-3’, 5’-GGTGCCTGATAACCTGAGTA-3’), TBK1 (5’-CATAAGCTTCCTTCGTCCAG-3’, 5’-ATCACTTCTTTATTCCTACG-3’), IRF3 (5’-GGGAGTGGGATTGTCCAAGC-3’, 5’-GGCACCAACAGCCGCTTCAG-3’). Lentiviral particles were harvested from supernatants of HEK293T cells that had been co-transfected with psPAX2 (Addgene 12260), pMD2.G (Addgene 12259), and the respective sgRNA-encoding derivative of lentiCRISPRv2. After filtration (0.45 µm), virus-containing supernatants were used to transduce A2EN cells in the presence of 8 µg/ml polybrene (Sigma-Aldrich). Cells were co-transduced with lentiviral vectors encoding the two distinct sgRNAs that target the same gene. Transduced cells were selected in presence of puromycin (1 µg/ml; Gibco) and cloned by limiting dilution.

### Confirmation of gene knockouts by western blot analysis

Protein extracts were prepared by cell lysis in boiling 1% SDS buffer, as previously described ^78^, separated by SDS PAGE (Mini-PROTEAN TGX 4-20% gels, Bio-Rad), and transferred onto nitrocellulose membranes (pore size of 0.2 µm, Bio-Rad). Membranes were then blocked for 20 minutes with 3% BSA in Tris-buffered saline with Tween (TBST; 20 mM Tris (pH 7.5), 150 mM NaCl, 0.1% Tween 20) and incubated overnight at 4°C with primary antibodies diluted in blocking buffer. The following primary antibodies were used: mouse-anti-β-actin (1:2000; Cell Signaling, 3700), rabbit-anti-ATG5 (1:1000; Cell Signaling 12994), rabbit-anti-ATG7 (1:1000; Cell Signaling, 8558), rabbit-anti-NDP52 (1:1000; Abcam ab68588); rabbit-anti-STING (1:1000; Cell Signaling, 13647), rabbit-anti-TBK1 (1:1000; Cell Signaling, 3504), and rabbit-anti-IRF3 (1:1000; Cell Signaling, 11904). After incubation, membranes were washed thrice with TBST, incubated for one hour with horse radish peroxidase (HRP)-conjugated secondary antibodies (anti-mouse: Biorad, 172- 1011; anti-rabbit: Thermo-Fisher-Scientific, G-21234; 1:10,000-1:50,000 diluted in blocking buffer), and washed again thrice with TBST. Membranes were then incubated for 1 min with HRP substrate (SuperSignal West Pico PLUS or SuperSignal West Atto, Thermo-Fisher-Scientific) and chemiluminescent signals were recorded with an Amersham Imager 680RGB (GE Healthcare). Membranes were stripped for 15-30 minutes with Restore Plus western blot stripping buffer (Thermo-Fisher-Scientific) and blocked with 3% BSA in TBST before detection of additional targets. Band intensities were quantified using Image Quant TL (GE Healthcare). Expression levels of target proteins were normalized to the expression of β-actin.

### Chlamydia strains

Infection experiments were carried out with the following *Chlamydia* strains: *C. trachomatis* L2/434/Bu (CTL2, ATCC VR-902B), an rsGFP-expressing derivative of CTL2 (CTL2-GFP; *i.e.*, CTL2 transformed with plasmid p2TK2-SW2-IncDProm-RSGFP-IncDTerm ^79^), *C. trachomatis* A/HAR-13 (DSMZ, 19440), *C. trachomatis* D/UW-3/Cx (DSMZ, 19411), *C. trachomatis* E/Bour (DSMZ, 19131), *C. caviae* GPIC ^80^, and *C. muridarum* MoPn (DSMZ, 28544). Two different procedures were used to prepare infection inocula, *i.e.*, density-gradient purified EBs (of CTL2 and CTL2-GFP) and crude preparations (of all other strains). Both procedures were described in detail in a previous publication ^78^. Bacterial preparations obtained by either procedure were resuspended in SPG (sucrose-phosphate-glutamate) buffer (75 g/l sucrose, 0.5 g/l KH_2_PO_4_, 1.2 g/l Na_2_HPO_4_, 0.72 g/l glutamic acid, pH 7.5), briefly sonicated, and stored at -80 °C. Bacteria were titered as previously described ^81^.

### Screening assay protocol

HeLa cells in suspension were mixed with CTL2-GFP (30 IFU/cell) and then centrifuged (800 x g, 5 min) to enhance infection. Subsequently, the infected cells were seeded (4,000 cells/well) in black transparent bottom 384-well plates (containing compounds or controls) using a Multidrop Combi liquid dispenser (Thermo-Fisher-Scientific) and the plates were incubated for 26 h (37 °C, 5% CO_2_). At that time, 1/5 volume of resazurin (Sigma-Aldrich; 0.15 mg/ml in Dulbecco’s phosphate-buffered saline (DPBS; Gibco)) was added followed by further incubation for 2.5 h. Resorufin and rsGFP fluorescence were measured at a Spark plate reader (Tecan) using wavelengths of 560/590 nm (excitation/emission) and 490/510 nm, respectively. Cells in plates intended for imaging were then fixed with 4% formaldehyde for 20 min at room temperature, washed two times with DPBS, and stored at 4 °C in DPBS until staining. Cells were stained with Hoechst 33342 (Invitrogen, 5 µg/ml in DPBS) for 15 min, washed three times with DPBS, and imaged at an ImageXpress Micro Confocal high-content imaging system (Molecular Devices), using the FITC filter for rsGFP fluorescence and the DAPI filter for Hoechst fluorescence. Image processing and detection of inclusions and host cell nuclei were performed with CellProfiler (version 4.0.7) ^82^. Total inclusion area in an image was calculated by multiplying the number of detected inclusions by the mean inclusion area in that image. Wells treated with ciprofloxacin (15 µg/ml) or staurosporine (1 µM) served as positive controls for complete bacterial growth inhibition and host cell toxicity, respectively. Wells treated solely with DMSO (vehicle control) served as negative control for both. Data was normalized to values obtained from DMSO-treated wells, after subtraction of background levels represented by the positive controls.

### Benchmarking the screening assay with antibiotics

Concentration series of chloramphenicol (0.005-500 µg/ml, Sigma-Aldrich), ciprofloxacin (0.003-300 µg/ml, Sigma-Aldrich), doxycycline (0.0002-20 µg/ml, Sigma-Aldrich), erythromycin (0.002-200 µg/ml, Sigma-Aldrich), gentamicin (0.01-1000 µg/ml, Thermo-Fisher-Scientific), kanamycin (0.01-1000 µg/ml, Duchefa), spectinomycin (0.02-2000 µg/ml, Sigma-Aldrich), and tetracycline (0.002-200 µg/ml, Molekula) were tested for inhibition of *C. trachomatis* growth and host cell toxicity using the bulk fluorescence readouts of the screening assay protocol (as described above). To mimic the conditions of the screen, the antibiotics were added to the plates before seeding the infected cells (*i.e.*, at 0 hpi). MICs were calculated using GraphPad Prism (version 8.4.3) based on a previously described method of data analysis ^83^.

### Compound library screen

The compound library screened experimentally was obtained from the Chemical Biology Consortium Sweden (CBCS), which owns a large collection of molecules (> 350,000) derived from various sources, in-house and commercial, and such donated by biotech companies. Specifically, we screened the CBCS primary screening set, which includes 36,785 small molecules selected to represent the diversity within the larger collection, but with a bias towards molecules with drug-like profiles. The compounds were obtained pre-dispensed in assay-ready black transparent bottom 384-well plates. Each compound was represented as a single well (final concentration after seeding of cells: 10 µM compound, 0.1% DMSO). Each plate also included the respective control wells, containing DMSO (final concentration 0.1%), ciprofloxacin (final concentration 15 µg/ml), or staurosporine (final concentration 1 µM). The compounds were tested for inhibition of *C. trachomatis* growth and host cell toxicity using the screening assay protocol (as described above). In the hit validation and potency determination steps, compounds were tested in duplicate and triplicate, respectively, with each replicate on separate plates. IC_50_ values were determined by fitting data into dose-response curves using the “[inhibitor] vs. response (three parameters)” equation in GraphPad Prism (version 8.4.3).

### Virtual screen and validations

Model development was based on the data from the experimental screen (all screened compounds, except for 380 molecules for which the structures were not available), supplemented with a set of compounds found to have been previously tested for antichlamydial activity according to a literature search. The literature search was conducted in PubMed and Web of Science, including studies published up until May 7^th^ 2020, using the following terms: “antichlamydial”, “novel *Chlamydia* therapy”, “*Chlamydia trachomatis* growth inhibition”, and “*Chlamydia trachomatis* inclusion inhibition”. For obvious reasons, only studies that included information on chemical structures were included. The literature compounds were binarized as hits or non-hits based on the same criterion as applied in the screen (*i.e.*, compounds reported to achieve > 50% bacterial growth inhibition at 10 µM were considered hits). It is worth noting that the literature compounds had been tested with different experimental methods and that their antichlamydial activities had been reported in various ways; usually either as IC_50_ or as percent inhibition at a single concentration. In the latter case, an estimated IC_50_ was calculated by extrapolating to 50% inhibition (if the reported inhibition was within 10-90%; otherwise the compound was excluded). Compounds with a molecular weight of over 1000 g/mol were also excluded. This left a total of 412 compounds. After curation of the expanded dataset (experimental and literature data), molecular descriptors (1D and 2D) were calculated for all included molecules by PaDEL-Descriptor (version 2.21) ^31^ with default settings, using as input SMILES strings to represent chemical structures. The molecular descriptors were then used to train a random forest classifier (default settings) with the Weka machine learning tool (version 3.8.4) ^84^. Prior to training the model, the synthetic minority oversampling technique (SMOTE), as implemented in Weka, was used to increase the number of instances in the minority (hit) class by a factor of three, and the majority (non-hit) class was undersampled to achieve a final ratio of 1:1 between the classes. The final model could classify an input compound as a hit or a non-hit and reported a prediction score of 0.5-1. The model was then applied to virtually screen the ChEMBL database ^34^ (release CHEMBL28) in Weka, using the same molecular descriptors (calculated by PaDEL-Descriptor) as in training. A total of 25 selected hit compounds were then purchased (from ChemBridge, TargetMol, or Vitas-M Laboratory) and experimentally tested for antichlamydial activity using the screening assay protocol described above, with bulk GFP fluorescence as the readout.

### Machine learning-aided identification of persistence inducers

To generate images for model training, HeLa cells infected with CTL2-GFP as in the screening assay described above were treated with concentration series of the beta-lactam antibiotics ampicillin (0.02- 2000 µg/ml, Duchefa), carbenicillin (0.02-2000 µg/ml, Duchefa), and penicillin G (0.002-200 U/ml, Sigma-Aldrich) and images of chlamydial inclusions were acquired as described above for the screening assay protocol. We chose to include rather high concentrations of beta-lactams to overcome the effects of the beta-lactamase expressed from the plasmid in CTL2-GFP. Image processing and detection of chlamydial inclusions were performed with CellProfiler (version 4.0.7) ^82^, with features extracted using the MeasureObjectSizeShape, MeasureObjectIntensity, MeasureObjectIntensityDistribution, and MeasureTexture measurement modules. These features were then used as input to train random forest and support vector machine (SMO) classifiers (default settings) with the Weka machine learning tool (version 3.8.4) ^84^. Images from the three highest concentrations of each antibiotic (with clear persistence induction) were used as ‘hits’ and images of DMSO-treated controls as well as of DMSO controls from the screen itself were used as ‘non-hits’. Before model training, SMOTE was used to increase the number of instances in the minority (hit) class to achieve a final ratio of 1:10 between the classes.

### Verification of persistence induction by confocal microscopy

Confluent HeLa cells, seeded at 150,000 cells/well in 8-well chamber slides (Ibidi), were infected with CTL2 (10 IFU/cell) and at the same time treated with compounds (10 µM). Positive control wells for persistence induction were treated with penicillin G (100 U/ml). The cells were incubated for 28 h (37 °C, 5% CO_2_), then fixed with 4% formaldehyde in DPBS for 20 min, and permeabilized for 15 min with 0.2% Triton X-100 in DPBS. Subsequently, the cells were incubated for 20 min in blocking solution (2% BSA in DPBS), and then for 1 hour with blocking solution containing primary antibodies (goat-anti-OmpA (1:1000; Thermo-Fisher- Scientific, PA1-7209) and rabbit-anti-Slc1 (1:500; ^85^)). Cells were then washed thrice with DPBS, incubated for 1 hour in blocking solution containing Hoechst 33342 (Invitrogen; 5 µg/ml) and secondary antibodies labeled with AlexaFluor 488 or AlexaFluor 555 (Invitrogen; 1:1000), and then washed again thrice with DPBS. Images were taken at a Leica SP8 confocal laser scanning microscope. Processing of confocal images and measurements of bacterial diameters were performed with LAS X Office (Leica, version 1.4.4).

### Verification of morphological recovery from persistence

HeLa cells in suspension were infected with CTL2-GFP (30 IFU/cell), centrifuged (800 x g, 5 min), and seeded (20,000 cells/well) in black transparent bottom 96-well plates (with compound-containing wells (10 µM) and control wells containing DMSO (0.2%) or penicillin G (100 U/ml)). The cells were then incubated for 28 h (37 °C, 5% CO_2_). At that time, the compounds were removed, and the cells were washed with DPBS, followed by addition of new medium. Cells to be imaged at 0 hours post-removal were immediately fixed with 4% formaldehyde as above. Other cells were further incubated for 24 and 48 hours, and then fixed. Prior to imaging, the cells were stained with Hoechst 33342 (Invitrogen; 5 µg/ml in DPBS) for 15 min and washed thrice with DPBS. Images were taken with an ImageXpress Micro Confocal high-content imaging system (Molecular Devices) as described above.

### Quantification of EB formation and its recovery from persistence

Quantification of EBs was performed essentially as previously described ^80^. Briefly, confluent HeLa cells in 96-well plates were infected with CTL2-GFP (5 IFU/cell) and at the same treated with compounds (at the indicated concentrations). At 40 hpi, cell lysates were prepared by incubation of the cells in sterile water, followed by addition of 1/4 volume of 5X SPG buffer. In experiments measuring recovery from persistence, compounds were washed off at 40 hpi by two washes with compound-free medium, followed by preparation of cell lysates at 40 hours post wash-off. To quantify the actual number of infectious particles (*i.e.*, EBs) in the initial inoculum (input) and the collected lysates (output), confluent monolayers of Vero cells in 96-well plates were infected with serial dilutions of the different samples. Inclusion numbers were determined by fluorescence microscopy at 28 hpi, as described above for the imaging of screening plates. Based on the number of inclusions found in the input and the output, the number of EBs formed in each infected cell could be determined. Values from wells containing medium only (blank) were subtracted, and the data was normalized to values from DMSO-treated control wells.

### Compound effects in different cell lines

The compounds were first diluted in medium containing CTL2-GFP (media containing 0.3% DMSO or 15 µg/ml ciprofloxacin were included as controls), and then transferred to confluent monolayers of cells in black transparent bottom 96-well plates. The infection dose used was cell-line-dependent: 5 IFU/cell for HeLa, Vero, JH4, and BALB/3T3, 10 IFU/cell for A2EN, and 25 IFU/cell for UMNSAH/DF-1. The plates were centrifuged (1500 x g, 30 min), and the cells were incubated for 26 h. At that time, 1/5 volume of resazurin (0.15 mg/ml in DPBS) was added followed by further incubation for 2.5 h. Resorufin and bulk GFP fluorescence were measured as described above. Compound effects in STING pathway- and autophagy-deficient A2EN cells were tested in the same way.

### Compound effects against different *Chlamydia* spp

Compound effects against different *Chlamydia* spp. were tested in a similar way as effects in different cell lines, but instead using different bacterial strains to infect HeLa cells only, with infection doses of 1.5 IFU/cell for CTL2-GFP, 0.5 IFU/cell for CTL2, 2 IFU/cell for CTA, 1 IFU/cell for CTD, CTE, and CC, and 0.3 IFU/cell for CM. After incubation, the cells were fixed and stained with an antibody targeting Slc1, as described above. Images were taken with an ImageXpress Micro Confocal high-content imaging system (Molecular Devices), and inclusion areas were measured with CellProfiler (version 4.0.7). Values from ciprofloxacin-treated wells or wells containing medium only (blank) were subtracted, and the data was normalized to values from DMSO-treated control wells. IC50 values were determined as described above.

### Compound effects on commensal bacteria in dilution assay

Cultures of *Escherichia coli* (DSM 301), *Lactobacillus reuteri* (DSM 20016), *Enterococcus faecalis* (DSM 20478), *Bacteroides thetaiotaomicron* (DSM 2079), and *Bifidobacterium adolescentis* (DSM 20083) were maintained in a Whitley H35 hypoxystation adapted to anaerobic conditions (37 °C, 0% O_2_, 5% CO_2_). Prior to their use in experiments, the bacteria were cultivated overnight in liquid Gifu Anaerobic Medium (GAM; HyServe), and then used to inoculate fresh cultures in GAM that were grown to exponential phase. In parallel, 96-well plates containing dilution series of the compounds to be tested were prepared under normoxic conditions in modified GAM (mGAM; HyServe). Azithromycin and doxycycline were included as controls for growth inhibition, and DMSO was included as vehicle control. The plates were then moved to the hypoxystation 3 h before inoculation with bacteria to allow the plates to equilibrate to anoxic conditions. Subsequently, the exponentially growing bacteria were diluted to an OD_600_ of 0.5 and then further 1:100 diluted, resulting in the final suspension used to inoculate the compound plates (typically 10^5^-10^6^ bacteria/well). The plates were then incubated for 18 h at 37 °C (0% O_2_, 5% CO_2_), followed by measurements of OD_600_ to estimate bacterial growth. Values from wells containing medium only (blank) were subtracted, and the data was normalized to values from DMSO-treated control wells.

### Compound effects on commensal bacteria in radial diffusion assay

Cultures of *Blautia coccoides* (DSM935), *Prevotella copri* (DSM18205), *Bacteroides fragilis* (DSM2151), *Dubosiella newyorkensis* (DSM103457), *Prevotella histicola* (DSM26979), *Enterococcus faecalis* (DSM 20478), *Bacteroides thetaiotaomicron* (DSM2079), and *Bifidobacterium adolescentis* (DSM20083) were incubated under anaerobic conditions (37 °C, 0% O_2_, 5% CO_2_) in a Whitley H35 hypoxystation. *Escherichia coli* (DSM 301) and *Lactobacillus reuteri* (DSM 20016) were grown aerobically at 37 °C. At the start of the assay, approximately 10_7_ CFU from mid-log cultures of the bacteria were mixed with 10 ml underlay gel consisting of 0.1% EEO-Agarose (Sigma), 0.1% broth powder [brain heart infusion broth (Merck Millipore) supplemented with 5 g/l yeast extract (Gibco) for *B. coccoides*, *B. fragilis*, and *B. thetaiotaomicron*; fastidious anaerobe broth (Neogen) for *P. copri*, *P. histicola*, and *D. newyorkensis*; tryptic soy broth (BD) for *E. coli* and *E. faecalis*; de Man, Rogosa and Sharpe broth (Merck Millipore) for *L. reuteri*; and reinforced clostridial medium (VWR) for *B. adolescentis*], and 10 mM sodium phosphate buffer (pH 7.4) and poured into an empty petri dish. 2 mm holes were punched into the underlay agar and 4 μl of 160 µM compound solution were added to the holes. After 3 hours of incubation, 10 ml of nutrient-rich overlay gel, consisting of 3% broth powder, 0.1% EEO-Agarose and 10 mM sodium phosphate buffer (pH 7.4), were added on top of the underlay gel. The petri dishes were incubated at 37 °C under aerobic (*E. coli* and *L. reuteri*) or anaerobic (all other bacteria) conditions for 24-48 hours. The results are reported as diameter of inhibition zone measured in mm.

### Antibiotic interactions

Black transparent bottom 96-well plates were prepared to contain known antibiotics and a selected set of our top compounds, each at its previously determined IC_50_ concentration. The antibiotics/compounds were added in pairwise combinations and single-antibiotic/compound wells were included for comparison. Ciprofloxacin (15 µg/ml) was used as control for complete bacterial growth inhibition and DMSO (0.1%) was used as vehicle control. HeLa cells in suspension were infected with CTL2-GFP (30 IFU/cell), centrifuged (800 x g, 5 min), and seeded in the compound-containing plates at 20,000 cells/well. The cells were then incubated for 28.5 h (37 °C, 5% CO2), followed by measurements of bulk GFP fluorescence. Data was normalized to values obtained from DMSO-treated wells, after subtraction of background levels represented by values from ciprofloxacin-treated wells. Interaction type (synergistic, antagonistic, or additive) was classified based on calculations of epistasis, as previously described ^42^.

### Compound effects on STING pathway- and autophagy activation

A2EN cells (wild-type) were seeded in 6-well plates at 1.5 x 10^5^ cells/well and incubated overnight. Cells were then either left uninfected or were infected with CTL2 (3 IFU/cell), followed by centrifugation (1500 x g, 30 min) and incubation (37 °C, 5% CO_2_). At 20 hpi, cells were treated with compounds (c1_e_ and c2_e_: 0.6 µM; c3_e_ and c5_e_: 3 µM; c4_e_: 4 µM) or a corresponding amount of DMSO. At 8 hours post treatment, protein samples were generated, and western blot analysis was conducted as described above. The following primary antibodies were used: mouse-anti-β-actin (1:2000; Cell Signaling, 3700); rabbit-anti-phospho-STING (1:1000; Cell Signaling, 19781), and rabbit-anti-LC3A/B (1:1000; Sigma-Aldrich, L8918). Band intensities were quantified using Image Quant TL (GE Healthcare). Expression levels of target proteins were normalized to the expression of β-actin. Values from compound-treated samples were then divided by values from DMSO-treated samples.

### Long-term treatment

HeLa cells in suspension were infected with CTL2-GFP (30 IFU/cell), centrifuged (800 x g, 5 min), and seeded in black transparent bottom 96-well plates at 20,000 cells/well, followed by incubation (37°C, 5% CO_2_). At 24 hpi, the medium was replaced with medium containing compounds or antibiotics (azithromycin and doxycycline). The cells were then further incubated, and the compounds were replenished at 48 and 72 hpi, followed by wash-off at 96 hpi. Finally, the cells were incubated until 168 hpi. Measurements of bulk GFP fluorescence were performed every 24 h. To visualize growth over time, data was normalized to values from ciprofloxacin-treated control wells (treated with ciprofloxacin (15 µg/ml) from 0 hpi, representing zero growth) at each time point. Images were also taken at each time point (or every 0.5 h during live cell imaging), using an ImageXpress Micro Confocal high-content imaging system (Molecular Devices). Long-term treatments in the presence of penicillin G (100 U/ml) were performed in a similar fashion.

### Statistics

Statistical analyses were performed with Graphpad Prism (version 8.4.3). Statistical significance is indicated as follows: *, P ≤ 0.05; **, P ≤ 0.01; ***, P ≤ 0.001; ****, P ≤ 0.0001. Principal component analysis and hierarchical clustering were done with SIMCA (version 17; Sartorius). Tanimoto coefficients (CDK standard) were calculated using a previously developed KNIME workflow ^86^.

### Availability of data and biological materials

This study includes no data deposited in external repositories. Data have been made available in the supplementary materials of the manuscript. Raw data files can be shared upon request. The authors are not able to provide compounds, but compound identifiers are listed in the supplementary tables and should allow purchase from commercial sources or compound libraries. Biological materials generated in this study can be shared upon request.

## Acknowledgements

The authors would like to acknowledge the support from the Chemical Biology Consortium Sweden (CBCS), its nodes at Umeå University and Karolinska Institutet, and in particular its staff Anna Eriksson and Åsa Slevin. CBCS is a Swedish national research infrastructure funded by the Swedish Research Council (VR- 2021-00179) and SciLifeLab. We would further like to thank the Umeå Hypoxia Research Facility and its manager Emilio Bueno for providing support and access to equipment for anaerobic cultivation of bacteria. Moreover, we acknowledge the Biochemical Imaging Center at Umeå University and the Swedish National Microscopy Infrastructure NMI (VR-2019-00217) for assistance with confocal microscopy. We are also grateful to Isabelle Dérre (University of Virginia) who had previously shared with us the plasmid enabling rsGFP expression in CTL2. This work was supported by grants from the Swedish Research Council (VR- 2016-06598, VR-2018-02286, VR-2021-06602, VR-2022-00852).

## Author contributions

Conceptualization, MÖ, BSS; Investigation, MÖ, DRV, KM, LM, JF, FPB, EC, MRD, KW, BSS; Formal analysis, MÖ, DRV, KM, LM, JF, FPB, EC, MRD, BSS; Visualization, MÖ; BSS; Writing – Original Draft, MÖ, BSS; Writing – Review & Editing, MÖ, KM, FPB, BOS, BSS; Supervision, BOS, BSS; Funding acquisition, BOS, BSS. All authors read and approved the final manuscript.

## Declaration of interests

The authors declare no competing interests.

## Supplementary information

Fig S1. Development of a screening assay for novel chemical inhibitors of *C. trachomatis* growth.

**(A)** Outline of the main steps and measurements in the screening assay. **(B)** Resorufin fluorescence at different HeLa cell seeding densities (mean ± SD, n = 3). The table shows Pearson’s r from a seeding density of 0 cells/well to the seeding density indicated. **(C)** GFP fluorescence at different infection doses (IFU/cell), with a seeding density of 4,000 cells/well (mean ± SD, n = 3, two-way ANOVA with Sidak’s multiple comparisons test of untreated vs. ciprofloxacin). **(D)** Host cell viability (via resorufin fluorescence) at different infection doses, with a seeding density of 4,000 cells/well (mean ± SD, n = 3, one-way ANOVA with Dunnett’s multiple comparisons test of infected vs. uninfected cells). **(E)** Representative images of HeLa cells infected in suspension with different amounts of CTL2-GFP. Since infection in suspension is less efficient than infection of adherent cells, even a dose of 30 IFU/cell left a significant number of cells uninfected. Scale bar is 10 µm. **(F)** DMSO tolerance of CTL2-GFP and HeLa cells, as measured by GFP and resorufin fluorescence, respectively (mean ± SD, n = 2 and n = 3, respectively, one-way ANOVA with Dunnett’s multiple comparisons test). **(G-H)** Plate uniformity and signal variability of the bacterial growth inhibition assay, based on measuring GFP fluorescence derived from CTL2-GFP (G), and the corresponding resazurin-based assay for host cell viability (H). Displayed are data from a representative of three independent experiments. In the left panels, each dot denotes a single well. NC, M, and PC refer to negative control, midlevel, and positive control for bacterial growth inhibition (G) and host cell toxicity (H), respectively.

**Fig S2. Assay performance benchmarked with clinical antibiotics. (A)** Eight clinical antibiotics were tested for bacterial growth inhibition and host cell viability with the bulk fluorescence readouts of the screening assay protocol (mean ± SD, n = 3). **(B)** MICs of eight antibiotics as determined in (A) (mean of n = 3) and compared with previously reported data. The calculated MIC of erythromycin is likely less accurate, due to the shape of its dose-response curve (see (A)). Beta-lactam antibiotics were not included in this analysis, as the plasmid driving GFP-expression in CTL2-GFP also encodes a beta-lactamase.

**Fig S3. Decision tree for classification of screening compounds as hits or non-hits.** The decision tree enabled integration of information from both bulk fluorescence measurements and high-content imaging for hit selection. Inclusion area was selected as the parameter of choice for image-based assessment of bacterial growth, as all compounds that would have been hits according to inclusion count were also hits according to inclusion area, but not vice versa.

**Fig S4. Chemical structures of the 52 priority compounds selected from the experimental compound library screen.** The structures were drawn based on their SMILES strings, using OpenBabel (version 3.0.0)

**Fig S5. Dose-response curves of the 52 priority compounds selected from the experimental compound library screen.** Data is shown for bulk GFP fluorescence as well as total inclusion area (mean ± SD, n = 3). The lines indicate the curve fits used for IC_50_ calculation, and the IC_50_ values are given in the respective plots.

**Fig S6. Dose-response curves of selected top compounds in different host cell lines. (A)** Growth inhibition by top compounds from the experimental compound library screen. The data is based on measurements of bulk GFP fluorescence and shows separate curves from three biological replicates. Error bars indicate standard deviations of three technical replicates. The lines indicate the curve fits used for IC_50_ calculation. **(B)** Growth inhibition by top compounds from the virtual screen, presented as in (A).

**Fig S7. Toxicity of selected top compounds against different host cell lines. (A)** Host cell viability after exposure to top compounds from the experimental compound library screen. The data is based on measurements of bulk resorufin fluorescence (mean ± SD, n = 3, two-way ANOVA with Dunnett’s multiple comparisons test of the lowest tested concentration of each compound vs. other concentrations). **(B)** Host cell viability after exposure to top compounds from the virtual screen, presented as in (A).

**Fig S8. Effect of top compounds on the growth of five species of gut bacteria.** The bacteria were grown in liquid medium containing compounds for 18 h, at which point OD_600_ was measured (mean ± SD, n = 3, two-way ANOVA with Sidak’s multiple comparisons test of each compound and concentration vs. the mean of all 0.03 µM samples for a particular species). Data was normalized to values obtained from DMSO-treated wells. AZM, azithromycin; DOX, doxycycline.

**Fig S9. Determination of between-compound interactions and interactions with clinical antibiotics.** Bacterial growth inhibition in HeLa cells infected with CTL2-GFP (30 IFU/cell, 28.5 hpi) and classification of interaction type for all pairwise combinations of fifteen selected top compounds with twelve clinical antibiotics (and c1_e-_c3_e_), tested at IC_50_ (mean ± SD, n = 3). Interaction type was classified based on calculations of epistasis, as previously described ^42^. Darker colors indicate stronger synergistic (green) or antagonistic (blue) interactions. Data for c1_e_-c5_e_ were also included in Fig 5B.

**Fig S10. Confirmation of gene knockouts by western blot analysis.** Western blot analysis of A2EN cell lines (wild-type (WT) or knockout (KO) for indicated genes) to confirm the absence of the targeted proteins in autophagy or the STING-pathway of the type I IFN response. Marked in green are the KO cell clones used in this study.

**Table S1. Host cell viability and *Chlamydia* growth data from the experimental compound library screen.** Data was normalized to values obtained from DMSO-treated wells, after subtraction of background levels.

**Table S2. The 271 hits from the experimental compound library screen retested at three concentrations.** Host cell viability and *Chlamydia* growth data were normalized to values obtained from DMSO-treated wells, after subtraction of background levels.

**Table S3. The 52 priority compounds from the experimental compound library screen, tested at eleven concentrations in triplicate.** Data was normalized to values obtained from DMSO-treated wells, after subtraction of background levels.

Table S4. Compilation of compounds previously tested for antichlamydial activity.

**Table S5. The 174 hits from the virtual screen remaining after a series of filtering steps.**

**Table S6. The 25 experimentally tested compounds from the virtual screen, tested at eleven concentrations in triplicate.** Data was normalized to values obtained from DMSO-treated wells, after subtraction of background levels.

## Notes

### Competing Interest Statement

The authors have declared no competing interest.

