## Supplementary Figures for "A multi-strategy antimicrobial discovery approach reveals new ways to combat *Chlamydia*"

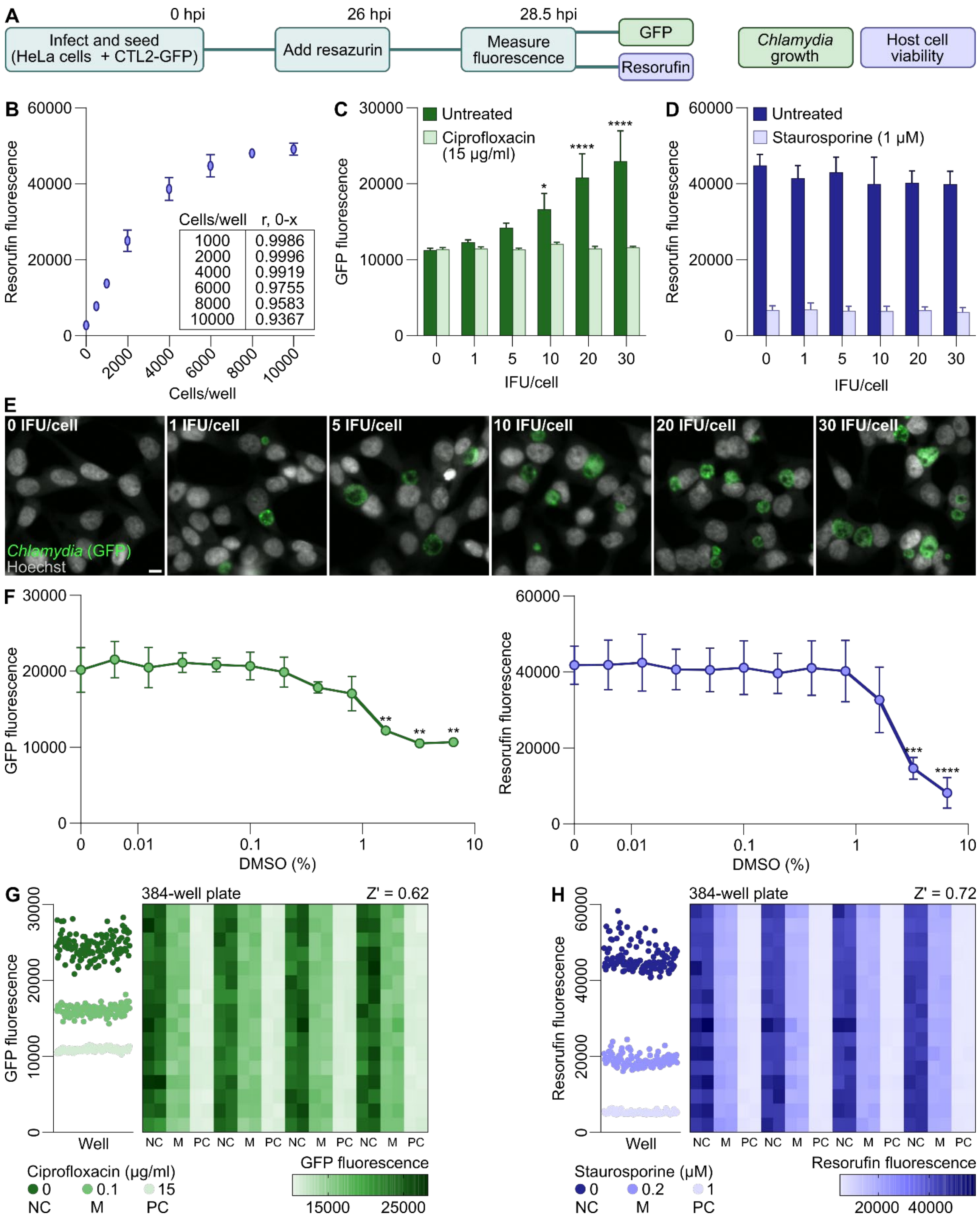
A
0 hpi
26 hpi
28.5 hpi
Infect and seed (HeLa cells + CTL2-GFP)
Add resazurin
Measure fluorescence
GFP
Resorufin
Chlamydia growth
Host cell viability
B
Resorufin fluorescence
Cells/well
Cells/well r, 0-x
1000 0.9986
2000 0.9996
4000 0.9919
6000 0.9755
8000 0.9583
10000 0.9367
C
GFP fluorescence
Untreated
Ciprofloxacin (15 µg/ml)
IFU/cell
D
Resorufin fluorescence
Untreated
Staurosporine (1 µM)
IFU/cell
E
0 IFU/cell
1 IFU/cell
5 IFU/cell
10 IFU/cell
20 IFU/cell
30 IFU/cell
Chlamydia (GFP)
Hoechst
F
GFP fluorescence
Resorufin fluorescence
DMSO (%)
G
GFP fluorescence
Well
384-well plate
Z' = 0.62
NC M PC NC M PC NC M PC NC M PC
Ciprofloxacin (µg/ml)
0 NC
0.1 M
15 PC
GFP fluorescence
15000 25000
H
Resorufin fluorescence
Well
384-well plate
Z' = 0.72
NC M PC NC M PC NC M PC NC M PC
Staurosporine (µM)
0 NC
0.2 M
1 PC
Resorufin fluorescence
20000 40000

**Fig S1. Development of a screening assay for novel chemical inhibitors of *C. trachomatis* growth. (A)** Outline of the main steps and measurements in the screening assay. **(B)** Resorufin fluorescence at different HeLa cell seeding densities (mean ± SD, n = 3). The table shows Pearson's r from a seeding density of 0 cells/well to the seeding density indicated. **(C)** GFP fluorescence at different infection doses (IFU/cell), with a seeding density of 4,000 cells/well (mean ± SD, n = 3, two-way ANOVA with Sidak's multiple comparisons test of untreated vs. ciprofloxacin). **(D)** Host cell viability (via resorufin fluorescence) at different infection doses, with a seeding density of 4,000 cells/well (mean ± SD, n = 3, one-way ANOVA with Dunnett's multiple comparisons test of infected vs. uninfected cells). **(E)** Representative images of HeLa cells infected in suspension with different amounts of CTL2-GFP. Since infection in suspension is less efficient than infection of adherent cells, even a dose of 30 IFU/cell left a significant number of cells uninfected. Scale bar is 10 µm. **(F)** DMSO tolerance of CTL2-GFP and HeLa cells, as measured by GFP and resorufin fluorescence, respectively (mean ± SD, n = 2 and n = 3, respectively, one-way ANOVA with Dunnett's multiple comparisons test). **(G-H)** Plate uniformity and signal variability of the bacterial growth inhibition assay, based on measuring GFP fluorescence derived from CTL2-GFP (G), and the corresponding resazurin-based assay for host cell viability (H). Displayed are data from a representative of three independent experiments. In the left panels, each dot denotes a single well. NC, M, and PC refer to negative control, midlevel, and positive control for bacterial growth inhibition (G) and host cell toxicity (H), respectively.

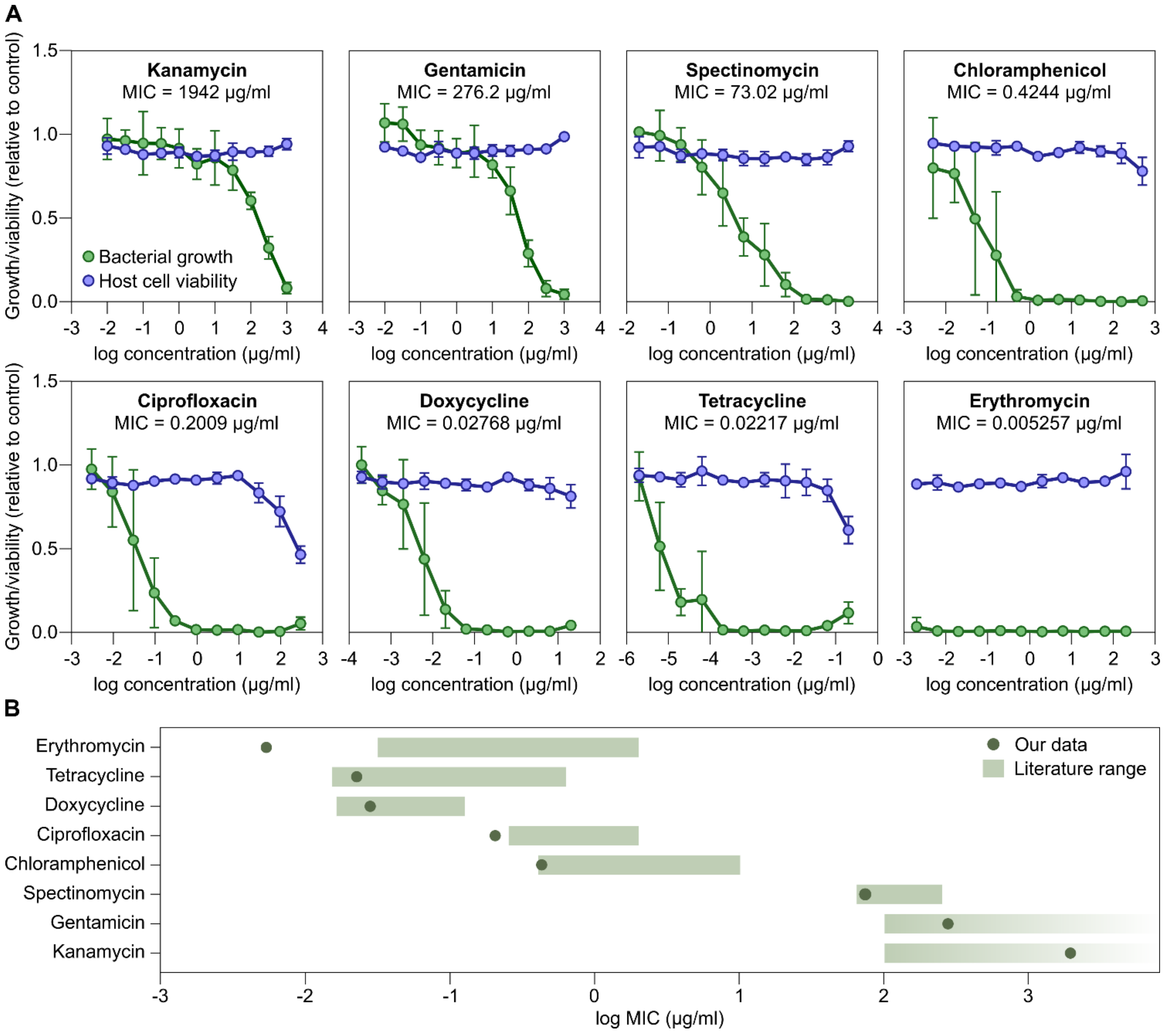


**Fig S2. Assay performance benchmarked with clinical antibiotics. (A)** Eight clinical antibiotics were tested for bacterial growth inhibition and host cell viability with the bulk fluorescence readouts of the screening assay protocol (mean ± SD, n = 3). **(B)** MICs of eight antibiotics as determined in (A) (mean of n = 3) and compared with previously reported data. The calculated MIC of erythromycin is likely less accurate, due to the shape of its dose-response curve (see (A)). Beta-lactam antibiotics were not included in this analysis, as the plasmid driving GFP-expression in CTL2-GFP also encodes a beta-lactamase.

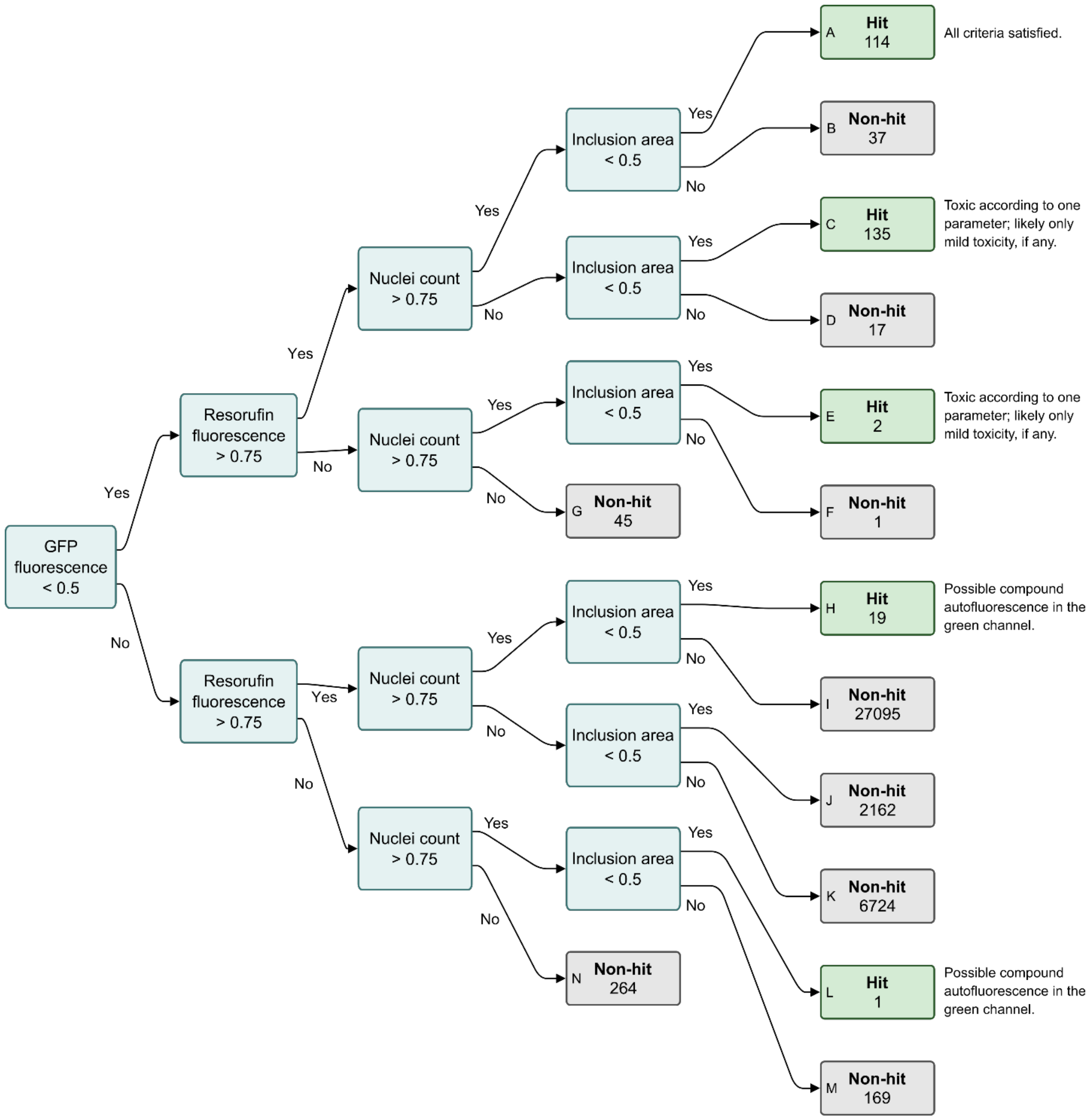


**Fig S3. Decision tree for classification of screening compounds as hits or non-hits.** The decision tree enabled integration of information from both bulk fluorescence measurements and high-content imaging for hit selection. Inclusion area was selected as the parameter of choice for image-based assessment of bacterial growth, as all compounds that would have been hits according to inclusion count were also hits according to inclusion area, but not vice versa.

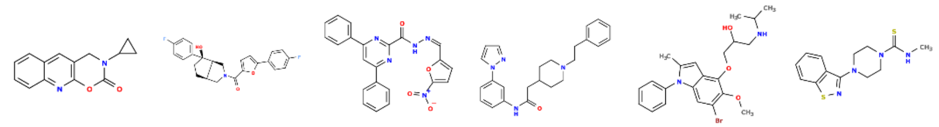

CBK282870 c1$_e$ | CBK286415 c2$_e$ | CBK261958 c3$_e$ | CBK285951 c4$_e$ | CBK263516 c5$_e$ | CBK288515 c6$_e$

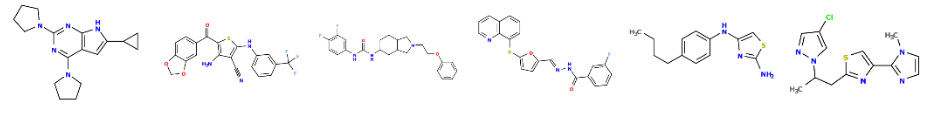

CBK016707G c7$_e$ | CBK074399 c8$_e$ | CBK083365T c9$_e$ | CBK263263 c10$_e$ | CBK020594 c11$_e$ | CBK304114 c12$_e$

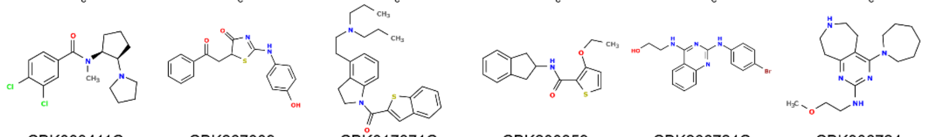

CBK003411C c13$_e$ | CBK267009 c14$_e$ | CBK017871C c15$_e$ | CBK200359 c16$_e$ | CBK266724C c17$_e$ | CBK306724 c18$_e$

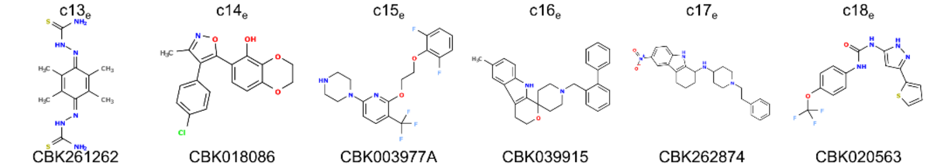

CBK261262 c19$_e$ | CBK018086 c20$_e$ | CBK003977A c21$_e$ | CBK039915 c22$_e$ | CBK262874 c23$_e$ | CBK020563 c24$_e$

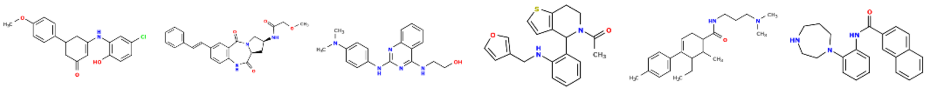

CBK263542 c25$_e$ | CBK289366 c26$_e$ | CBK268392C c27$_e$ | CBK286432 c28$_e$ | CBK016514 c29$_e$ | CBK038927T c30$_e$

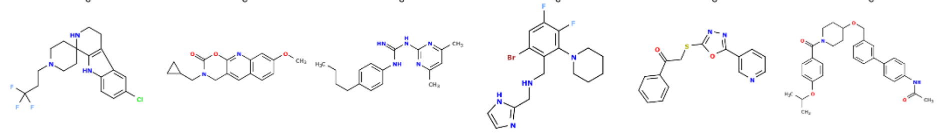

CBK039331 c31$_e$ | CBK282432 c32$_e$ | CBK262967 c33$_e$ | CBK304243 c34$_e$ | CBK068446 c35$_e$ | CBK149485 c36$_e$

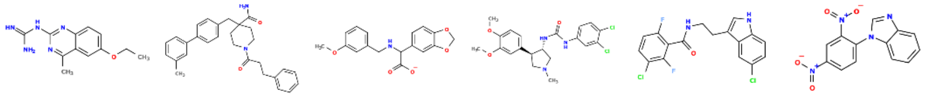

CBK010384C c37$_e$ | CBK285796 c38$_e$ | CBK015420D c39$_e$ | CBK081111 c40$_e$ | CBK159756 c41$_e$ | CBK261120 c42$_e$

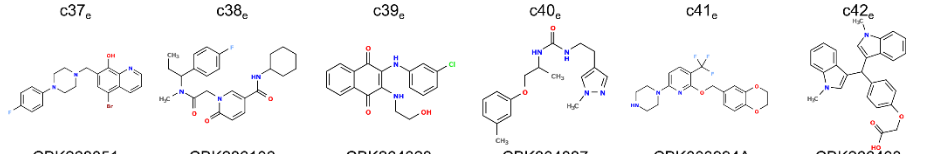

CBK268051 c43$_e$ | CBK286106 c44$_e$ | CBK264323 c45$_e$ | CBK304887 c46$_e$ | CBK003994A c47$_e$ | CBK266403 c48$_e$

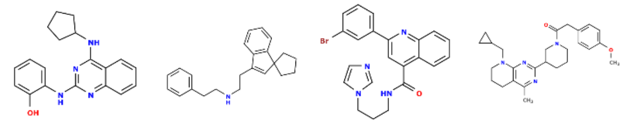

CBK268402C c49$_e$ | CBK024723 c50$_e$ | CBK265476 c51$_e$ | CBK286228 c52$_e$

**Fig S4. Chemical structures of the 52 priority compounds selected from the experimental compound library screen.** The structures were drawn based on their SMILES strings, using OpenBabel (version 3.0.0) [1].

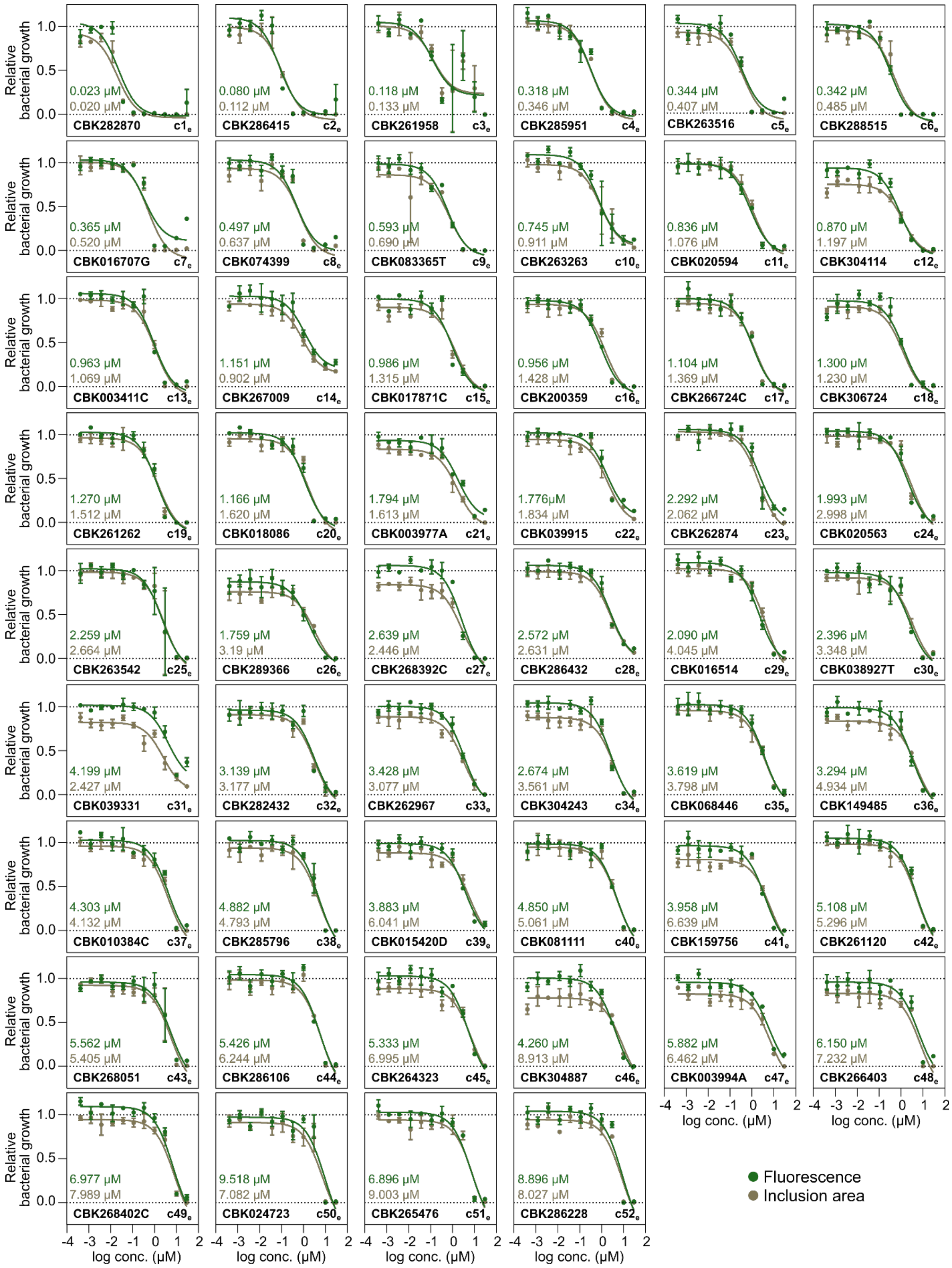
Relative bacterial growth
log conc. (µM)
Fluorescence
Inclusion area
0.023 µM 0.020 µM CBK282870 c1e
0.080 µM 0.112 µM CBK286415 c2e
0.118 µM 0.133 µM CBK261958 c3e
0.318 µM 0.346 µM CBK285951 c4e
0.344 µM 0.407 µM CBK263516 c5e
0.342 µM 0.485 µM CBK288515 c6e
0.365 µM 0.520 µM CBK016707G c7e
0.497 µM 0.637 µM CBK074399 c8e
0.593 µM 0.690 µM CBK083365T c9e
0.745 µM 0.911 µM CBK263263 c10e
0.836 µM 1.076 µM CBK020594 c11e
0.870 µM 1.197 µM CBK304114 c12e
0.963 µM 1.069 µM CBK003411C c13e
1.151 µM 0.902 µM CBK267009 c14e
0.986 µM 1.315 µM CBK017871C c15e
0.956 µM 1.428 µM CBK200359 c16e
1.104 µM 1.369 µM CBK266724C c17e
1.300 µM 1.230 µM CBK306724 c18e
1.270 µM 1.512 µM CBK261262 c19e
1.166 µM 1.620 µM CBK018086 c20e
1.794 µM 1.613 µM CBK003977A c21e
1.776µM 1.834 µM CBK039915 c22e
2.292 µM 2.062 µM CBK262874 c23e
1.993 µM 2.998 µM CBK020563 c24e
2.259 µM 2.664 µM CBK263542 c25e
1.759 µM 3.19 µM CBK289366 c26e
2.639 µM 2.446 µM CBK268392C c27e
2.572 µM 2.631 µM CBK286432 c28e
2.090 µM 4.045 µM CBK016514 c29e
2.396 µM 3.348 µM CBK038927T c30e
4.199 µM 2.427 µM CBK039331 c31e
3.139 µM 3.177 µM CBK282432 c32e
3.428 µM 3.077 µM CBK262967 c33e
2.674 µM 3.561 µM CBK304243 c34e
3.619 µM 3.798 µM CBK068446 c35e
3.294 µM 4.934 µM CBK149485 c36e
4.303 µM 4.132 µM CBK010384C c37e
4.882 µM 4.793 µM CBK285796 c38e
3.883 µM 6.041 µM CBK015420D c39e
4.850 µM 5.061 µM CBK081111 c40e
3.958 µM 6.639 µM CBK159756 c41e
5.108 µM 5.296 µM CBK261120 c42e
5.562 µM 5.405 µM CBK268051 c43e
5.426 µM 6.244 µM CBK286106 c44e
5.333 µM 6.995 µM CBK264323 c45e
4.260 µM 8.913 µM CBK304887 c46e
5.882 µM 6.462 µM CBK003994A c47e
6.150 µM 7.232 µM CBK266403 c48e
6.977 µM 7.989 µM CBK268402C c49e
9.518 µM 7.082 µM CBK024723 c50e
6.896 µM 9.003 µM CBK265476 c51e
8.896 µM 8.027 µM CBK286228 c52e

**Fig S5. Dose-response curves of the 52 priority compounds selected from the experimental compound library screen.** Data is shown for bulk GFP fluorescence as well as total inclusion area (mean ± SD, n = 3). The lines indicate the curve fits used for $IC_{50}$ calculation, and the $IC_{50}$ values are given in the respective plots.

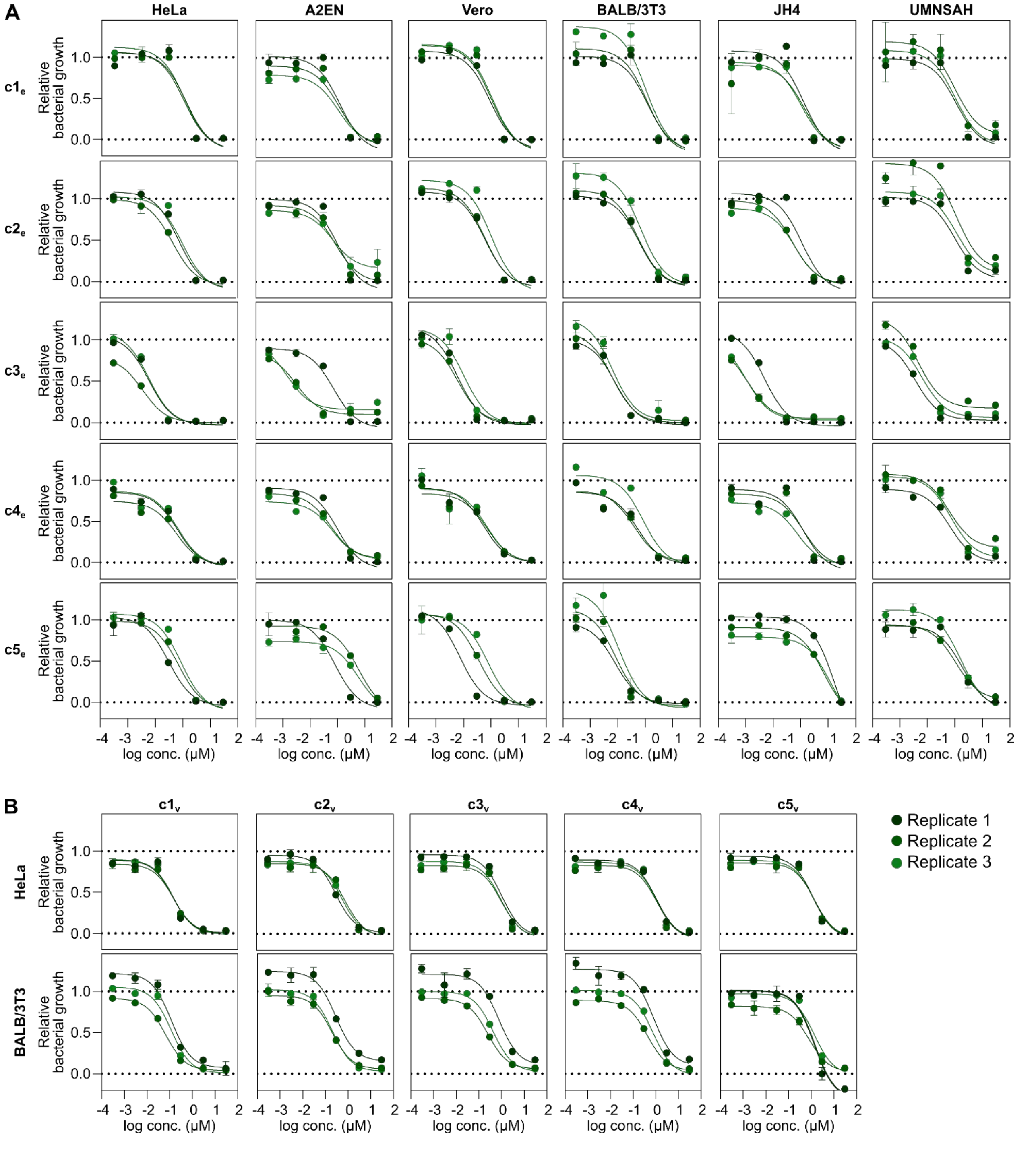


**Fig S6. Dose-response curves of selected top compounds in different host cell lines. (A)** Growth inhibition by top compounds from the experimental compound library screen. The data is based on measurements of bulk GFP fluorescence and shows separate curves from three biological replicates. Error bars indicate standard deviations of three technical replicates. The lines indicate the curve fits used for $IC_{50}$ calculation. **(B)** Growth inhibition by top compounds from the virtual screen, presented as in (A).

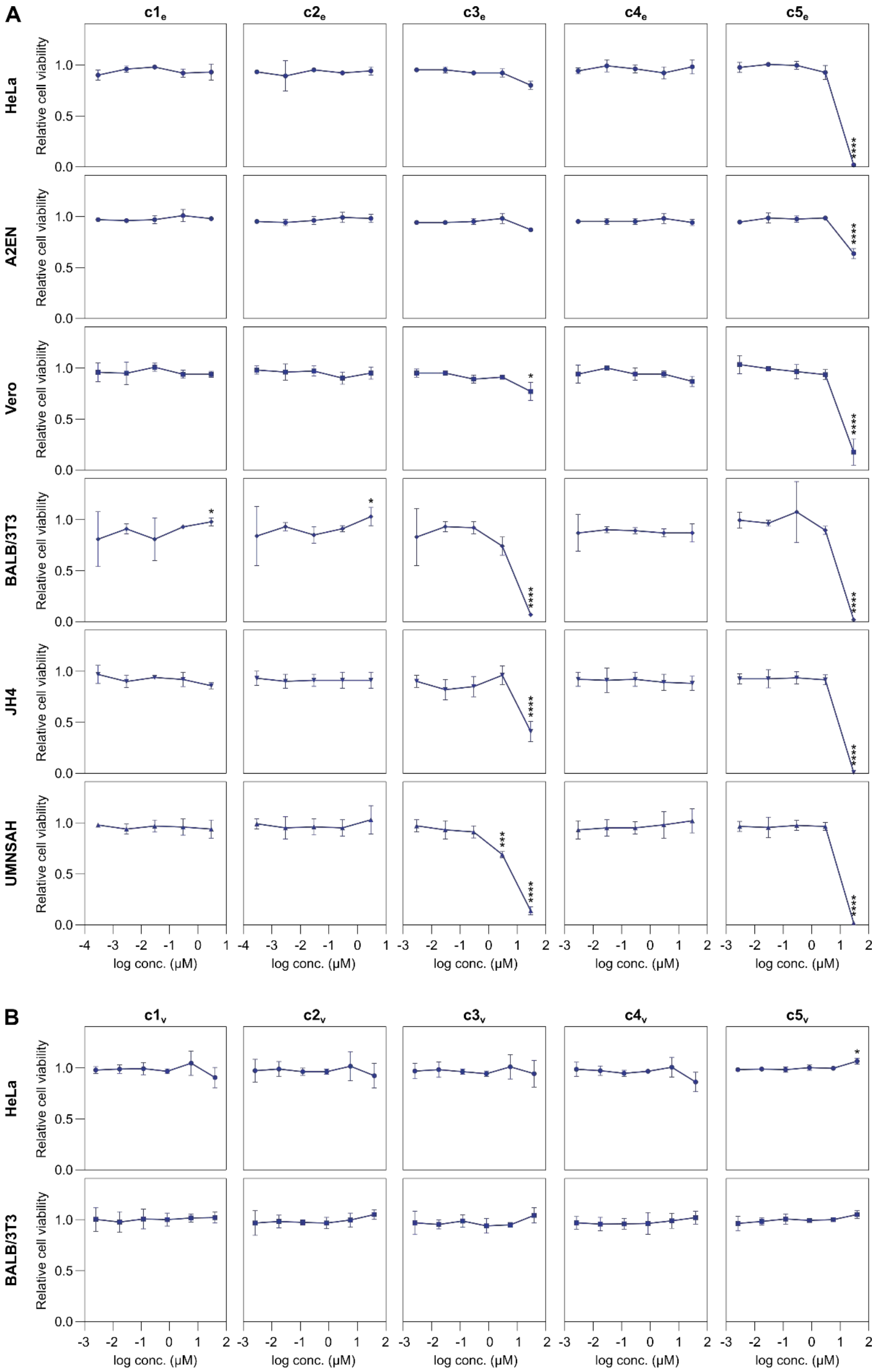
A
B
$c1_e$
$c2_e$
$c3_e$
$c4_e$
$c5_e$
$c1_v$
$c2_v$
$c3_v$
$c4_v$
$c5_v$
HeLa
A2EN
Vero
BALB/3T3
JH4
UMNSAH
Relative cell viability
log conc. (μM)

**Fig S7. Toxicity of selected top compounds against different host cell lines. (A)** Host cell viability after exposure to top compounds from the experimental compound library screen. The data is based on measurements of bulk resorufin fluorescence (mean ± SD, n = 3, two-way ANOVA with Dunnett's multiple comparisons test of the lowest tested concentration of each compound vs. other concentrations). **(B)** Host cell viability after exposure to top compounds from the virtual screen, presented as in (A).

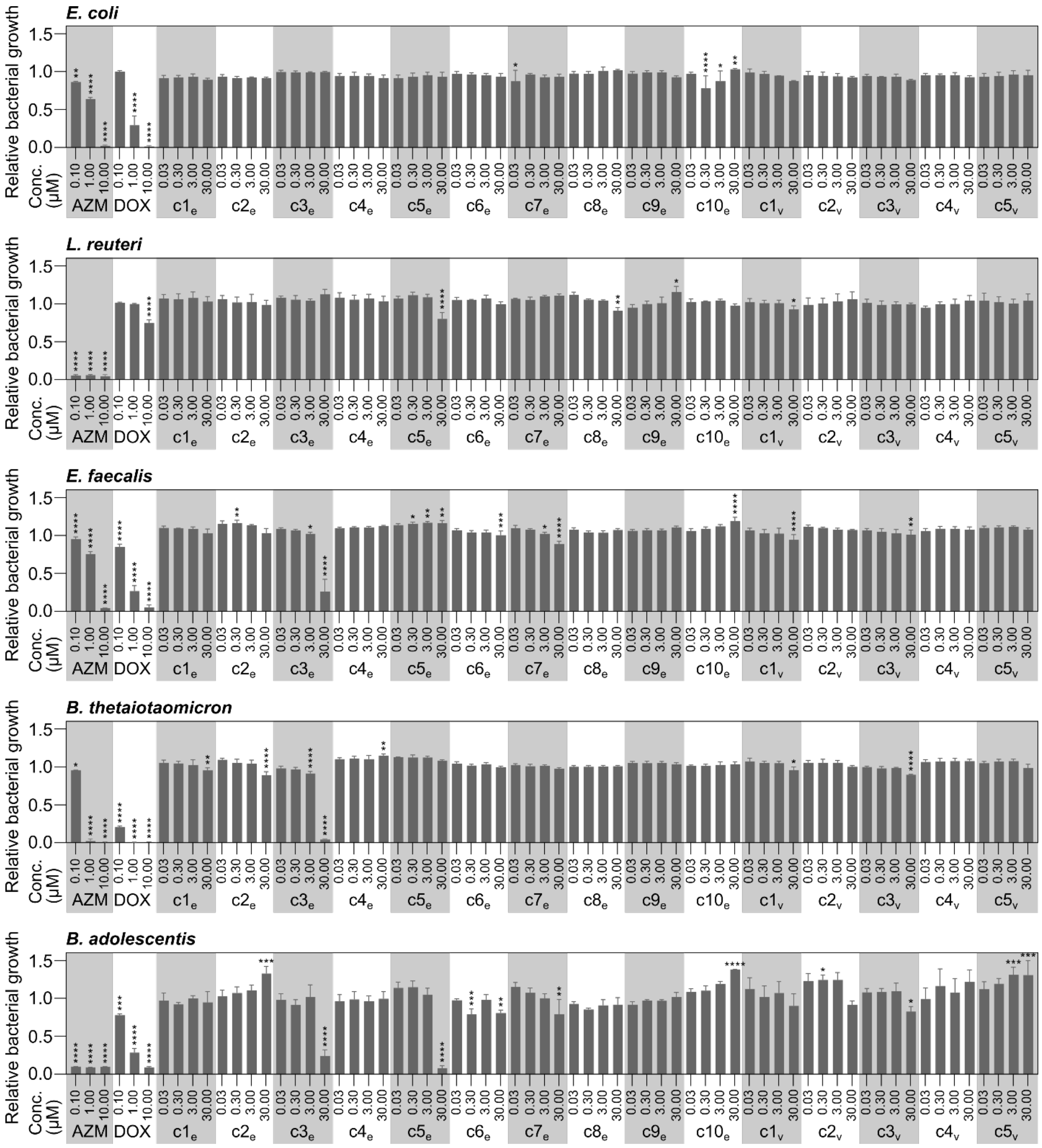


**Fig S8. Effect of top compounds on the growth of five species of gut bacteria.** The bacteria were grown in liquid medium containing compounds for 18 h, at which point $OD_{600}$ was measured (mean ± SD, n = 3, two-way ANOVA with Sidak's multiple comparisons test of each compound and concentration vs. the mean of all 0.03 μM samples for a particular species). Data was normalized to values obtained from DMSO-treated wells. AZM, azithromycin; DOX, doxycycline.

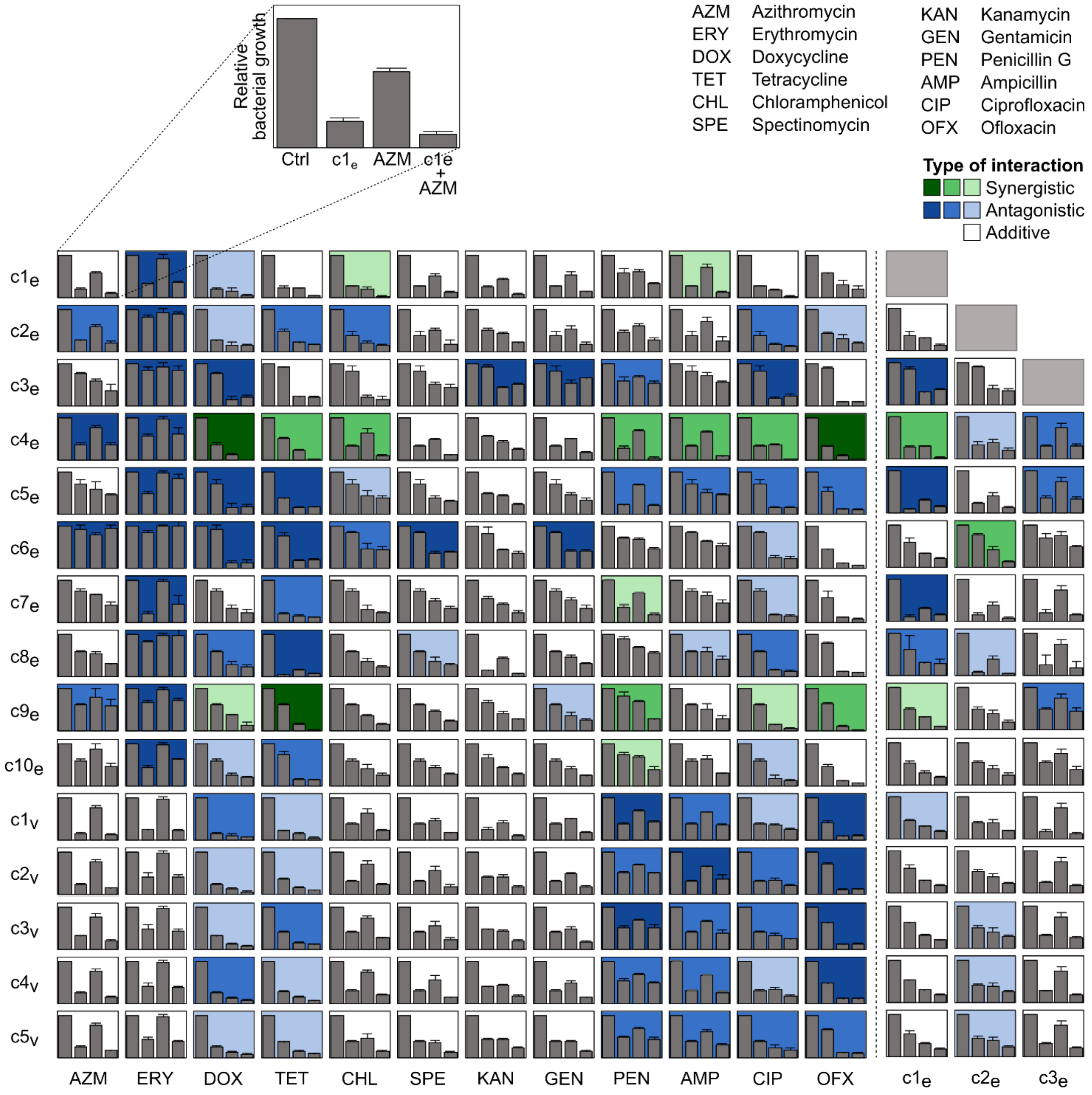


**Fig S9. Determination of between-compound interactions and interactions with clinical antibiotics.** Bacterial growth inhibition in HeLa cells infected with CTL2-GFP (30 IFU/cell, 28.5 hpi) and classification of interaction type for all pairwise combinations of fifteen selected top compounds with twelve clinical antibiotics (and $c1_e$-$c3_e$), tested at $IC_{50}$ (mean ± SD, n = 3). Interaction type was classified based on calculations of epistasis, as previously described [2]. Darker colors indicate stronger synergistic (green) or antagonistic (blue) interactions. Data for $c1_e$-$c5_e$ were also included in Fig 5B.

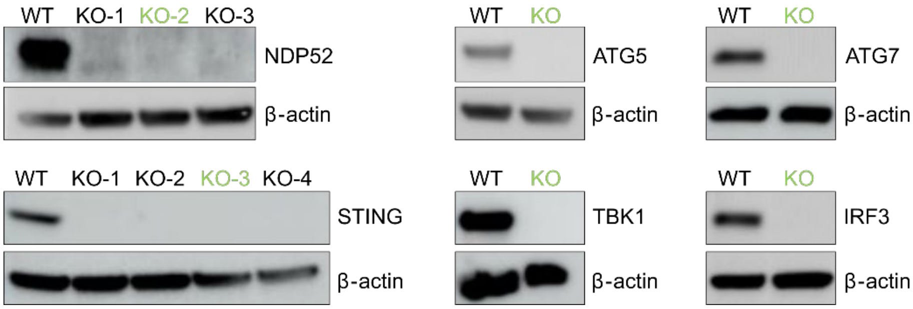


**Fig S10. Confirmation of gene knockouts by western blot analysis.** Western blot analysis of A2EN cell lines (wild-type (WT) or knockout (KO) for indicated genes) to confirm the absence of the targeted proteins in autophagy or the STING-pathway of the type I IFN response. Marked in green are the KO cell clones used in this study.
